# Structural evolution of an immune evasion determinant shapes Lyme borreliae host tropism

**DOI:** 10.1101/2022.09.13.507797

**Authors:** Ashley L. Marcinkiewicz, Kalvis Brangulis, Alan P. Dupuis, Thomas M. Hart, Maxime Zamba-Campero, Tristan A. Nowak, Jessica L. Stout, Inara Akopjana, Andris Kazaks, Janis Bogans, Alexander T. Ciota, Peter Kraiczy, Sergios-Orestis Kolokotronis, Yi-Pin Lin

**Author notes:** correspondence: Kalvis Brangulis, Ph.D., Latvian Biomedical Research and Study Centre, Riga, Latvia LV-1067; Sergios-Orestis Kolokotronis, Ph.D., Department of Epidemiology and Biostatistics. School of Public Health SUNY Downstate Health Sciences University, 450 Clarkson Avenue, MSC43A, Brooklyn, NY 11203, USA; Yi-Pin Lin, Ph.D., Wadsworth Center, NYSDOH, Albany, NY 12208. These authors have contributed equally to this work (Ashley L. Marcinkiewicz, Kalvis Brangulis). Yale University School of Medicine, New Haven, CT.

## Abstract

The preferential adaptation of pathogens to specific hosts, known as host tropism, evolves through host-pathogen interactions. Transmitted by ticks and maintained primarily in rodents and birds, the Lyme disease-causing bacterium *Borrelia burgdorferi* (*Bb*) is an ideal model to investigate the mechanisms of host tropism. In order to survive in hosts and escape complement-mediated clearance, a first-line host immune defense, *Bb* produces the outer surface protein CspZ that binds to the complement inhibitor factor H (FH) to facilitate bacterial dissemination in vertebrates. Despite high sequence conservation, CspZ variants vary in human FH-binding ability. Together with the FH polymorphisms found amongst vertebrate hosts, these findings raise a hypothesis that minor sequence variation in a bacterial outer surface protein confers dramatic differences in host-specific, FH-binding-mediated infectivity. We tested this hypothesis by determining the crystal structure of the CspZ-human FH complex, identifying a minor change localized in the FH-binding interface, and uncovered that the bird and rodent FH-specific binding activity of different CspZ variants directly impacts infectivity. Swapping the divergent loop region in the FH-binding interface between rodent- and bird-associated CspZ variants alters the ability to promote rodent- and bird-specific early-onset dissemination. By employing phylogenetic tree thinking, we correlated these loops and respective host-specific, complement-dependent phenotypes with distinct CspZ lineages and elucidated evolutionary mechanisms driving CspZ emergence. Our multidisciplinary work provides mechanistic insights into how a single, short pathogen protein motif could greatly impact host tropism.

**AUTHOR SUMMARY:** Lyme disease presents a suitable model for the investigation of host tropism – a pathogen’s ability to colonize and survive in different host species – since its causative agent, the spirochete *Borrelia burgdorferi* (*Bb*) is transmitted by ticks and maintained in rodent and bird reservoir hosts. In order to survive in vertebrates and escape from killing by complement, a first-line host immune defense, *Bb* produces the outer surface protein CspZ that binds the complement inhibitor factor H (FH) to promote infection. Protein sequence conservation seems to be linked to FH-binding activity divergence, raising the hypothesis that even minor variation can confer host-specific, FH-binding-mediated infectivity. Our work shows that that this minor variation is located in a loop in the CspZ protein localized in the CspZ-FH binding interface. Our functional experiments prove that this loop promotes bird- or rodent-specific FH-binding activity and infectivity. Swapping loops between rodent- and bird-associated CspZ variants alters their capability to confer host-specific dissemination. We further investigated the evolutionary mechanisms driving the emergence of the CspZ loop-mediated, host-dependent complement evasion. This multifaceted work demonstrates how a single, short protein motif can significantly impact host tropism.

## INTRODUCTION

The emergence of most infectious disease outbreaks often involves changes in host tropism, the preferential adaptation of pathogens to selectively invade and persist in hosts (1, 2). Such host tropism is often the result of ongoing host-pathogen interactions (3). Evolution theoretically favors the emergence of host-specializing pathogens, but host-generalist strategies can be advantageous in environments when pathogens have the potential to regularly interact with multiple hosts (4). Specialism vs. generalism can be attributed to polymorphisms within pathogen proteins that differentially interact with host ligands (3). Such polymorphism-mediated host ranges are often thought to require complex host-specific adaptive mechanisms (e.g., (5, 6)) that can be conferred by merely a few amino acids (7–9). Understanding how such minor differences impact diverse host-adapted phenotypes can elucidate the mechanistic insights for the emergence of modern infectious diseases, allowing the development of earlier and more efficient public health interventions.

Species within the *Borrelia* (also known as *Borreliella*) *burgdorferi* sensu lato genospecies complex cause Lyme disease. These spirochetal bacteria are transmitted by ticks and maintained by several reservoir hosts, primarily rodents and birds (10, 11). Lyme borreliae have been genotyped using different polymorphic loci, such as *ospA*, *ospC*, the 16S-23S r<u>R</u>NA intergenic <u>s</u>pacer (RST[yping]), and the loci included in <u>m</u>ultilocus <u>s</u>equence typing (MLST) (12, 13). Laboratory and field studies have shown that not only *Borrelia* species differ in the host species they infect, but that individual genotypes within single spirochete species display distinct preferential host associations, particularly within *B. burgdorferi* sensu stricto (hereafter *B. burgdorferi*), the primary Lyme disease agent in North America (14, 15). This species- and genotype -specific host selectivity distinguishes Lyme borreliae as a model system for the study of the molecular basis and evolutionary history of host tropism (14, 15).

Lyme borreliae host associations require the spirochetes to transmit from infected ticks to hosts, establish an infection at tick biting sites, and disseminate hematogenously to persist at distal tissues (14, 15). The survival of Lyme borreliae during these discrete events necessitate evasion of multiple host immune responses, including complement, a first-line innate defense mechanism of vertebrate extracellular fluid and blood (16–18). This powerful immune mechanism can be activated by the classical, lectin, or alternative pathways (19). The activation of these pathways results in the cascading cleavage and recombination of multiple complement components, ultimately resulting in phagocytic clearance, inflammation, and pore formation on the pathogen surface by the C5b-9 protein complex to lyse the cells (20). Complement is downregulated by diverse regulatory proteins to inhibit activation, preventing native cell damage in the absence of pathogens (21). For example, the alternative pathway is inhibited by factor H (FH), which is comprised of 20 individual short consensus repeat (SCR) domains (22, 23).

Similar to many bloodborne pathogens that evolved to circumvent complement-mediated clearance (20), Lyme borreliae produce several complement-inhibitory outer surface proteins (16–18). These proteins bind and recruit complement components and/or regulatory proteins on spirochete surface to inactivate complement (16–18). One of these proteins is a FH-binding protein, CspA (also known as <u>C</u>omplement <u>R</u>egulator <u>A</u>cquiring <u>S</u>urface <u>P</u>rotein 1, CRASP-1) that we demonstrated to confer survival of spirochetes via complement evasion in the tick bloodmeal (24). However, this protein is downregulated after spirochetes infect hosts suggesting a possibilities of other functionally redundant proteins to facilitate spirochete survival at this infection stage. In fact, CspZ (also known as CRASP-2), which binds to the 6^th^ and 7^th^ domains of SCR from FH (SCR6-7), is upregulated when spirochetes reside in hosts (25, 26). This protein is highly conserved both between and within different Lyme borreliae species (>85% amino acid (aa) identity between species and 98% between *B. burgdorferi* strains) (27–30). We and others have previously reported that FH-binding activity of one CspZ variant promotes spirochete survival in vertebrate sera, and consequently promotes host infectivity when non-physiologically relevant infection routes are used (i.e., not tick-transmitted) (31, 32). However, CspZ variants differ in their human FH-binding activity (27, 28), and vertebrate complement components or regulatory proteins vary between host taxa (e.g., ∼40% aa identity between mammalian and avian FH) (33). These findings raise several intriguing questions: Could such a minor divergence among CspZ variants confer host-specific differences in FH-binding activity, resulting in varying host infectivity phenotypes? Further, if those minor divergences are indeed a determinant of host tropism, how did such divergences evolve to impact Lyme borreliae-host associations?

In this study, we solved the high-resolution structure of the CspZ-human FH complex, identified a polymorphic CspZ motif within the FH-binding interface, and defined its contribution to host-specific FH-binding activity. We then examined the role of this motif in dictating strain-specific, host-dependent dissemination using mice and quail as rodent and avian models, respectively. Paired with evolutionary analyses of *cspZ*, we further elucidated how the evolutionary history behind a minor divergence in an immune evasion determinant can impact host tropism.

## RESULTS

### A polymorphic CspZ loop structure defines human FH-binding activity

We set to identify the regions driving the differences in human FH-binding activity among CspZ variants. We first pinpointed the amino acids that are involved in FH-binding by co-crystalizing the SCR6-7 of human FH and a CspZ variant with human FH-binding activity from *B. burgdorferi* strain B408 (CspZ_B408_) (28). The resulting crystal structure at 2.59Å showed an extensive binding interface between CspZ_B408_ and both SCR6 and SCR7 (**Fig. 1A**). Specifically, Asp47, Tyr50, Asn51, Thr54, Asn58, and Thr62 in helix-B, Arg142 in helix-F, Asn183 from helix-G, and Tyr214 in helix-I of CspZ_B408_ interacted with SCR7 (**Fig. 1B, Table S1**). Additionally, Asp71, from the loop between helix-B and -C, and Asp73 and Ser75 from the same region, bound to SCR7 and SCR6, respectively (**Fig. 1B, Table S1**).

**Figure 1.**
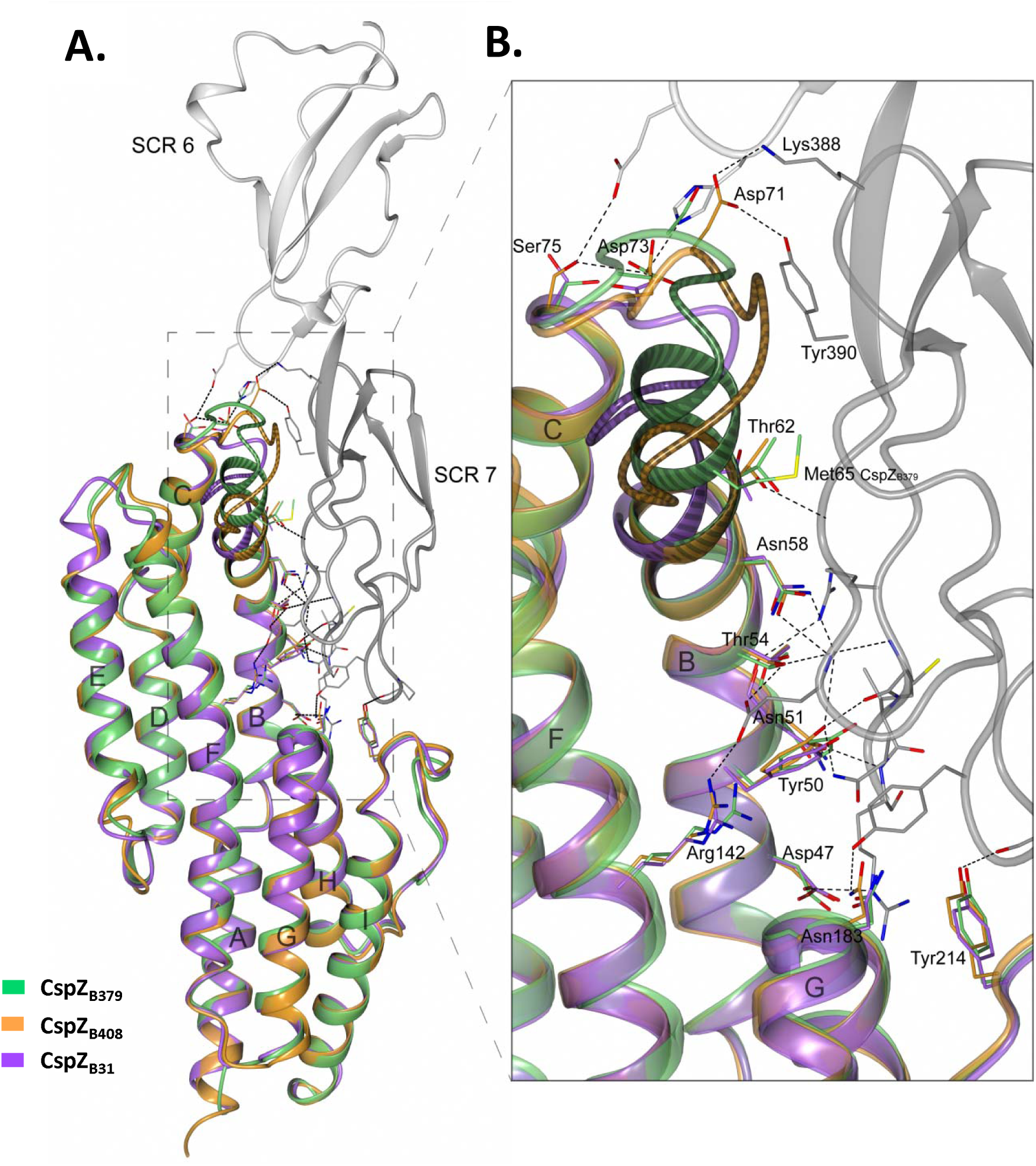
Superimposed crystal structures of CspZ_B408_-SCR6-7 complex, CspZ_B31_, and CspZ_B379_ reveals the FH-binding mechanisms of CspZ variants. (**A**) The crystal structure of human FH domains SCR6 (light grey) and SCR7 (dark grey) in complex with CspZ_B408_ (orange) was superimposed with that of CspZ_B31_ (purple) and CspZ_B379_ (green). (**B**) The FH-binding interface with the location of the CspZ_B408_ residues involved in human SCR6-7 binding, and the equvalent residues in CspZ_B31_ and CspZ_B379_. The numbering of the residues is given for CspZ_B408_. α-helices in CspZ are labelled from A to I starting from the N-terminus. The polymorphic loop region between α-helices B and C in CspZ_B408_ (residues 60-IMTYSEVNNVTD-71), CspZ_B31_ (residues 60-IMTYSEGT-67) and CspZ_B379_ (residues 60-IMTYIMTYSEGT-71) are indicated.

To locate the residues that impact CspZ FH-binding activity, we aligned the sequences of CspZ_B408_ with another CspZ variants that binds to human FH (i.e., CspZ from *B. burgdorferi* B31, CspZ_B31_), as well as CspZ that lacks human FH-binding ability from strain B379 (CspZ_B379_)(28). All the aforementioned residues involved in human FH-binding are conserved among all three variants, except Asn51 and Asp71 (**Fig. S1A**). Such conservation also reflects to the three-dimensional overlayed structure of the previously resolved CspZ_B31_ (30) with our newly-resolved crystal structures of CspZ_B379_ and CspZ_B408_ (2.10 and 2.45Å, respectively) (**Fig. 1A)**. For those two non-conserved amino acids, Asn51 of CspZ_B408_ is substituted to serine in CspZ_B379_, but the superimposed structure suggests this would not inhibit human FH-binding in CspZ_B379_ (**Fig. 1B**). Asp71 of CspZ_B408_ interacts with Lys388 and Tyr390 of SCR6, but this residue is part of an insertion that is not present in CspZ_B31_ and CspZ_B379_ (**Fig. 1B**). Therefore, these non-conserved residues alone cannot explain the differential human FH binding ability of these CspZ variants. However, the C-terminal helix-B and the following loop region between helix-B and -C do harbor polymorphisms (**Fig. 1A)**, with the unique duplication of the last residues in helix-B (Ile60, Met61, Thr62, and Tyr63) of CspZ_B379_ (**Fig. S1A**), resulting in an extended helix-B in this variant (**Fig. 1B**). Such a structural extension leads to steric hinderance between CspZ_B379_ and human FH to selectively prevent human FH-binding activity by CspZ_B379_. Overall, our results reveal variant-specific CspZ structural differences in a loop and the adjacent helix-B (hereafter, “loop structures”), providing mechanistic insights into the CspZ-mediated, polymorphic human FH-binding activity.

### CspZ loop structures determine host-specific FH-binding activity

Similar to CspZ, SCR6-7 is divergent among vertebrate species (**Fig. S1B**), suggesting a role of the CspZ loop structures in dictating host-specific FH-binding activity. We found that CspZ_B31_ and CspZ_B408_ –but not CspZ_B379_– bind to mouse FH, whereas CspZ_B31_ and CspZ_B379_ –but not CspZ_B408_– bind to quail FH (**Fig. 2A and S2, Table S2**). We then searched for structural evidence of this FH-binding activity by superimposing the complex structure of CspZ_B408_-human FH with CspZ_B31_, CspZ_B379_, as well as the previously resolved structures of mouse SCR6-7, and the AlphaFold-predicted structure of quail SCR6-7 (pLDDT > 70) (34, 35) (**Fig. 2C**). We noted the similar tertiary structures between mouse and human SCR6-7, consistent with the ability of FH from both origins selectively binds to CspZ_B31_ and CspZ_B408_ (28) (**Table S2**). Similar to human FH, the CspZ_B379_-specific duplication by the extended C-terminal helix-B showed potential structural hinderance in the mouse FH-binding interface (**Fig. 2D**). However, compared to a loop region within human and mouse SCR7, the equivalent region in quail SCR7 (median pLDDT: 92) is positioned away from the CspZ-FH-binding interface, allowing sufficient space for extended helix-B of CspZ_B379_ to interact with quail FH (**Fig. 2D**). Further, the crystal structure of CspZ_B408_-SCR6-7 superimposed with the predicted structure of quail SCR6-7 showed that Asp71 within the loop of CspZ_B408_ may collide with the opposite loop region of quail FH, while the equivalent region in mouse SCR7 would not interfere with the binding of CspZ_B408_ (**Fig. 2D**).

**Figure 2.**
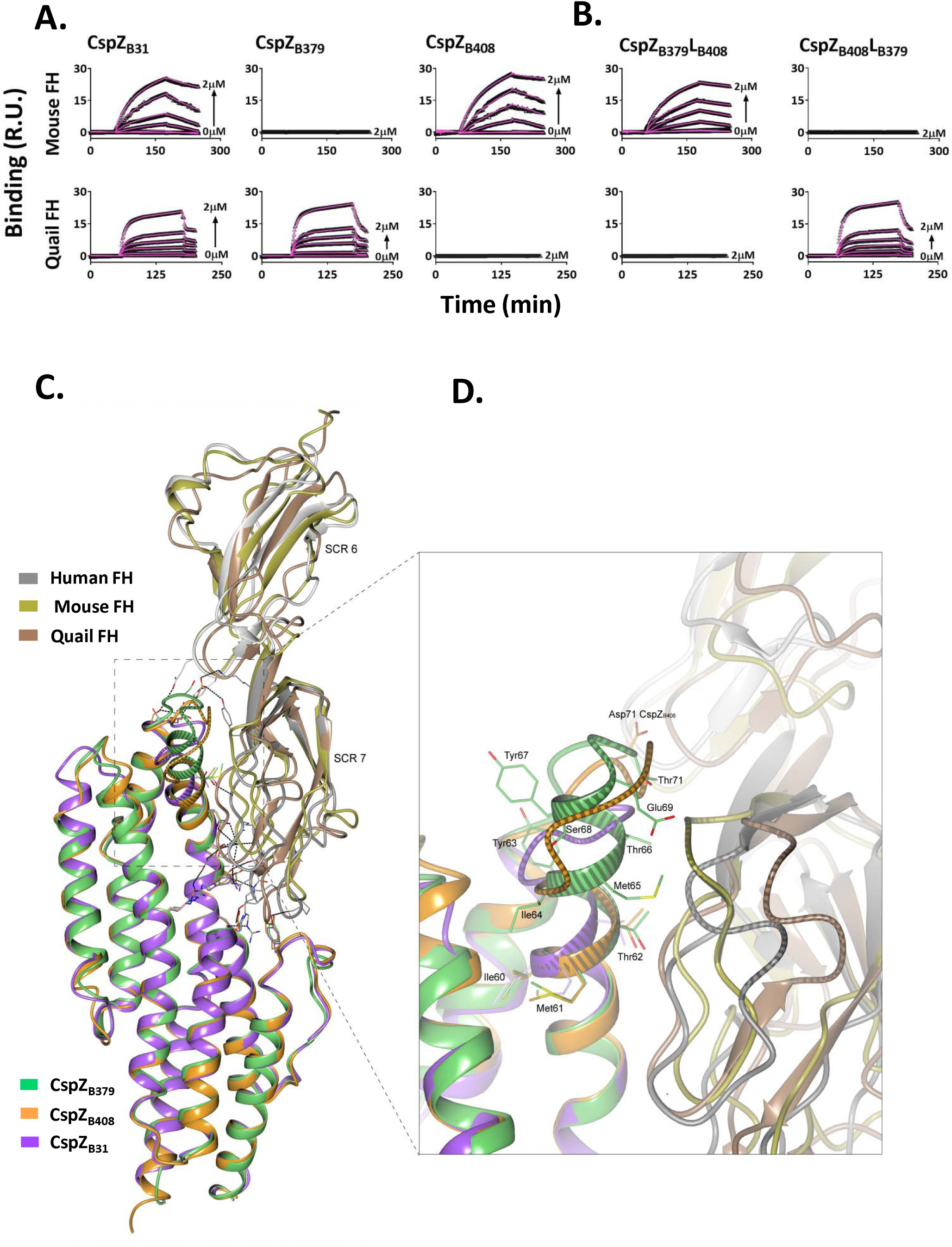
The polymorphic loop region in CspZ promotes host-specific FH-binding ability. **(A, B)** Ten micrograms of mouse or quail FH were conjugated on a SPR chip. 0.008 to 2μM of untagged **(A)** CspZ_B31_, CspZ_B379_, CspZ_B408_, **(B)** CspZ_B379_L_B408_, or CspZ_B408_L_B379_ was flowed over the chip surface. Binding was measured in response units (RU) by SPR. The experiments were performed with a single preparation of recombinant proteins tested in three independent replicates with samples ran in duplicate. The k_on_, k_off_, and K_D_ values were determined from the average of these three experiments (**Table 1**). Shown is one representative experiment. (**C**) The crystal structure of CspZ_B408_-SCR6-7 where human SCR6-7 (grey) is superimposed with mouse SCR6-7 (gold, PDB: 2YBY), and the predicted structure of quail SCR6-7 (brown), while CspZ_B408_ (orange) is superimposed with CspZ_B31_ (purple) and CspZ_B379_ (green). (**D**) The loop region in CspZ proteins located between α-helices B and C. The Asp71 in CspZ_B408_, and residues 60-IMTYIMTYSEGT- 71 in CspZ_B379_ are highlighted. The polymorphic loop region between α-helices B and C in CspZ variants, as well as the loop region in human, mouse and quail SCR6-7 located at the interface with helix-B in CspZ, is showed with a striped pattern.

These structural differences among FH from mouse/human vs. quail, pairing with the respective impacting AAs from different CspZ variants, suggest a possibility of the loop structures of CspZ to determine CspZ allelically different, host-specific FH-binding activity. To test this possibility, we swapped the loop regions of the two CspZ variants with distinct host-specific FH-binding ability to generate two chimeric proteins: CspZ_B379_L_B408_ has the backbone of a quail FH-binder (CspZ_B379_) and the loop structures of a mouse FH-binder (CspZ_B408_), whereas CspZ_B408_L_B379_ has the backbone of CspZ_B408_ and the loop structures of CspZ_B379_ (**Fig. S1A**). We found that CspZ_B379_L_B408_ and CspZ_B408_L_B379_ selectively bound to mouse and quail FH, respectively, demonstrating the CspZ loop structures are a determinant of host-specific FH-binding activity (**Fig. 2B and S2**).

### The CspZ loop structures dictate spirochete strain- and host-specific complement inactivation

We next examined the host-dependent complement inactivation of these CspZ variants produced on the spirochete surface. We thus obtained a wild-type *B. burgdorferi* strain B31-A3 (WT B31-A3) and a *cspZ*-deficient mutant strain in this background harboring an empty vector (Δ*cspZ*). We complemented this mutant with a plasmid encoding *cspZ_B31_* (pCspZ_B31_) (32), *cspZ_B379_* (pCspZ_B379_), or *cspZ_B408_* (pCspZ_B408_), or one of the two loop-swapped mutants (pCspZ_B379_L_B408_ and pCspZ_B408_L_B379_). We verified there was a similar generation time of these strains (**Table S3A**), and that there were no differences in the CspZ surface production level (**Fig. S3B**). Additionally, surface-produced CspZ bound FH in the same host-specific manner as the recombinant proteins (**Fig. S4**). We then determined the ability of each of these CspZ variants and mutants to inactivate mouse and quail complement by measuring the deposition levels of mouse C5b-9 and quail C8, respectively, in the presence of sera from each animal using flow cytometry (**Fig. 3A-B**). All strains had levels of mouse C5b-9 or quail C8 deposition significantly lower than the high passage, non-infectious, and mouse and quail complement-susceptible control strain *B. burgdorferi* B313 (**Fig. 3C-D**)(26). The Δ*cspZ* strain had significantly greater levels of mouse C5b-9 and quail C8 deposition than WT B31-A3 or pCspZ_B31_ (**Fig. 3C-D**)(32, 36). Compared to Δ*cspZ*, pCspZ_B379_ had indistinguishable levels of mouse C5b-9 deposition but significantly lower levels of quail C8, whereas pCspZ_B408_ harbored significantly lower levels of mouse C5b-9 but indistinguishable levels of quail C8 (**Fig. 3C-D**). Further, pCspZ_B379_L_B408_ recruited significantly lower levels of mouse C5b-9 but indistinguishable levels of quail C8 than Δ*cspZ*, while pCspZ_B408_L_B379_ bound indistinguishable levels of mouse C5b-9 but significantly lower levels of quail C8 than Δ*cspZ* (**Fig. 3C-D**). These results correspond with these variants’ host-specific FH-binding activity, suggesting that the CspZ loop structures-driven host-specific FH-binding activity confers host-specific complement inactivation.

**Figure 3.**
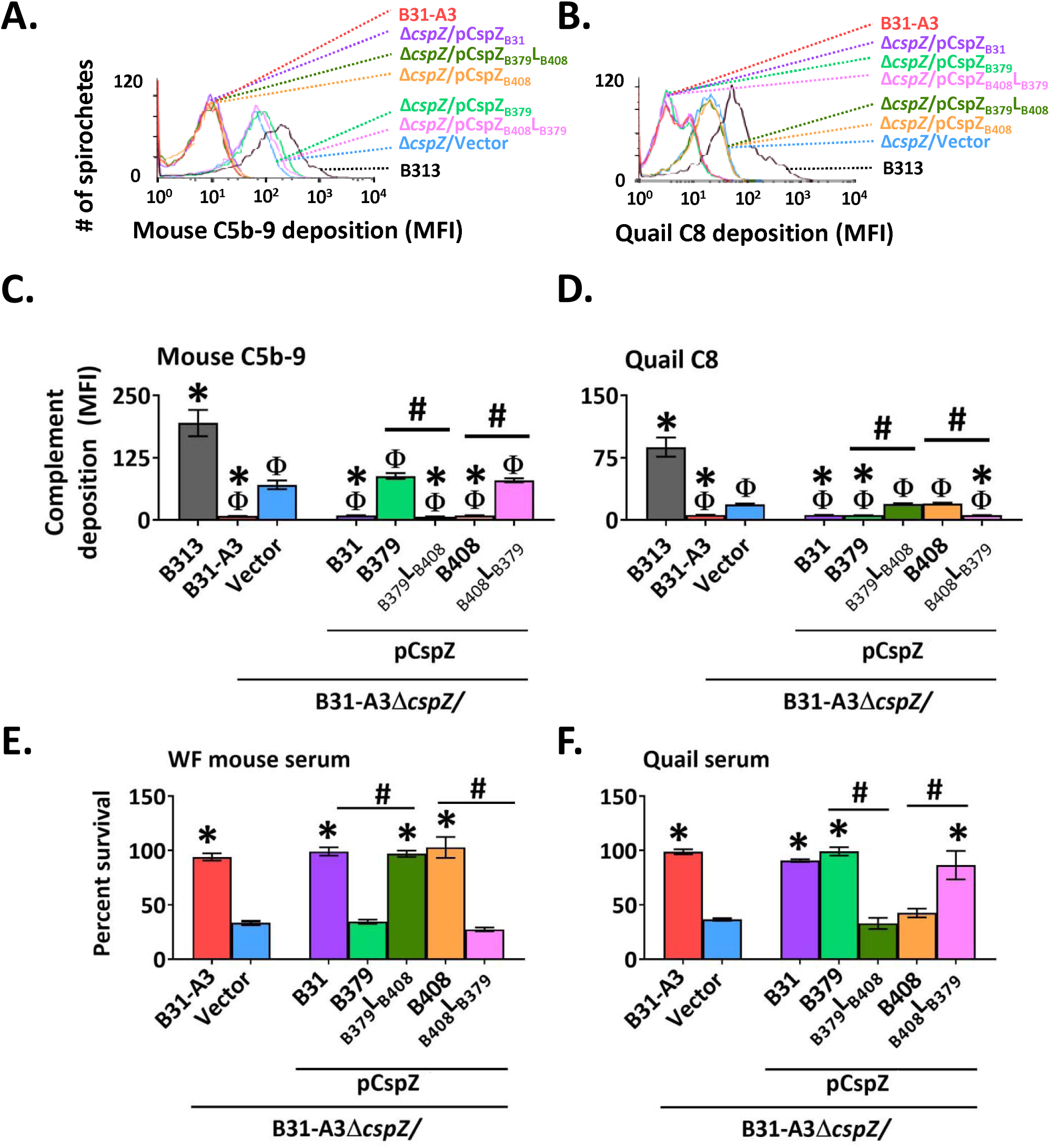
The CspZ loop-driven, variable FH-binding activity confers host-specific complement inactivation on spirochete surface and serum resistance. (**A to D**) *B. burgdorferi* strains B313 (negative control), B31-A3, B31-A3Δ*cspZ* harboring the empty vector pKFSS (“Δ*cspZ*/Vector”), or this *cspZ* mutant producing CspZ_B31_, CspZ_B379_, CspZ_B379_L_B408_, CspA_B408_, or CspZ_B408_L_B379_, were incubated with mouse or quail sera at a final concentration of 20%. The bacteria were stained with antibodies that recognize mouse C5b-9 or quail C8 prior to analysis by flow cytometry. Shown are the representative histograms of the analysis presenting the deposition levels of **(A)** mouse C5b-9 or **(B)** quail C8 on the surface of the indicated *B. burgdorferi* strains. The deposition levels of **(C)** mouse C5b-9 or **(D)** quail C8 on the surface of the indicated strains were measured by flow cytometry and presented as mean fluorescence index (MFI). Each bar represents the mean of three independent experiments ± SEM. Significant differences (p *<* 0.05, Kruskal-Wallis test with the two-stage step-up method of Benjamini, Krieger, and Yekutieli) in the deposition levels of mouse C5b-9 or quail C8 relative to B313 (“Φ”), Δ*cspZ*/Vector (“*”), or between two strains relative to each other (“#”), are indicated. (**E, F**) The *B. burgdorferi* strains were incubated for 4-h with (**E)** white-footed mouse sera or **(F)** quail sera, to a final concentration of 40%. The number of motile spirochetes was assessed microscopically. The precent survival of the strains was calculated using the number of motile spirochetes at 4-h post incubation normalized to that prior to incubation with sera. Each bar represents the mean of three independent experiments ± SEM. Significant differences (p *<* 0.05, Kruskal-Wallis test with the two-stage step-up method of Benjamini, Krieger, and Yekutieli) in the percent survival of spirochetes relative to the Δ*cspZ*/Vector (“*”) or between two strains relative to each other (“#”) are indicated.

We also measured the survival of these strains in rodent and quail sera. Note that sera from white-footed mice (*Peromyscus leucopus*), rather than house mice (*Mus musculus)*, were used to represent rodent sera, as complement of house mouse is labile *in vitro*, leading to inconsistent results (37, 38). Both WT B31-A3 and pCspZ_B31_ survived in white-footed mouse and quail sera more efficiently than Δ*cspZ* (**Fig. 3E-F**) (32). Compared to Δ*cspZ*, pCspZ_B379_ survived at significantly greater levels in quail but not white-footed mouse sera, whereas pCspZ_B408_ survived at significantly higher levels in white-footed mouse but not quail sera (**Fig. 3E-F**). Relative to Δ*cspZ*, pCspZ_B379_L_B408_ survived significantly higher in white-footed mouse but not in quail sera, whereas pCspZ_B408_L_B379_ survived significantly higher in quail but not in white-footed mouse sera (**Fig. 3E-F**). However, these differences were not observed in the presence of sera treated with Cobra Venom factor (CVF) or *Ornithodorus moubata* complement inhibitor (OmCI) that deplete functional rodent and quail complement, respectively (**Fig. S5**) (39, 40), suggesting that CspZ loop structures determine spirochete host-specific serum survival.

### The CspZ loop structures define host-specific, complement-dependent, early onset dissemination

We further evaluated if the CspZ loop structure variants determine host infectivity by allowing *Ixodes scapularis* nymphs carrying similar loads of each of the aforementioned *B. burgdorferi* strains to feed on mice and quail (**Fig. S6A, S6D**). The burdens of Δ*cspZ* were significantly lower than WT B31-A3 at 7- and 10-days post feeding (dpf) in mouse blood, and at 10 dpf in distal tissues (joints, heart, and bladder), and at 9 dpf in quail blood and distal tissues (brain and heart) (**Fig. S7**). There were no differences in burdens by a later time point (i.e., 14 dpf, **Fig. S7**). We thus performed subsequent experiments on 10 and 9 dpf in mouse and quail, respectively, for the other of *cspZ*-complemented strains. In mice, we found indistinguishable spirochete burdens at initial infection sites (tick bite sites) between all seven strains tested at 10dpf (**Fig. 4A**). The strains pCspZ_B31_, pCspZ_B408_, and pCspZ_B379_L_B408_, but neither pCspZ_B379_ nor pCspZ_B408_L_B379_, colonized mouse blood and distal tissues at significantly higher levels than Δ*cspZ* (**Fig. 4B-E**). At 9dpf in quail, we found no significantly different spirochete burdens at tick bite sites among strains. However, pCspZ_B31_, pCspZ_B379_, and pCspZ_B408_L_B379_, but neither pCspZ_B408_ nor pCspZ_B379_L_B408_, colonized quail blood and distal tissues at significantly greater levels than Δ*cspZ* (**Fig. 4F-I**). There were no significantly different burdens of any strain in any tissues from mice and quail with complement deficiency (C3^-/-^ mice or OmCI-treated quail) (**Fig. S8**). These results demonstrate that the CspZ loop structures define complement-dependent, spirochete strain- and host-specific early onset dissemination.

**Figure 4.**
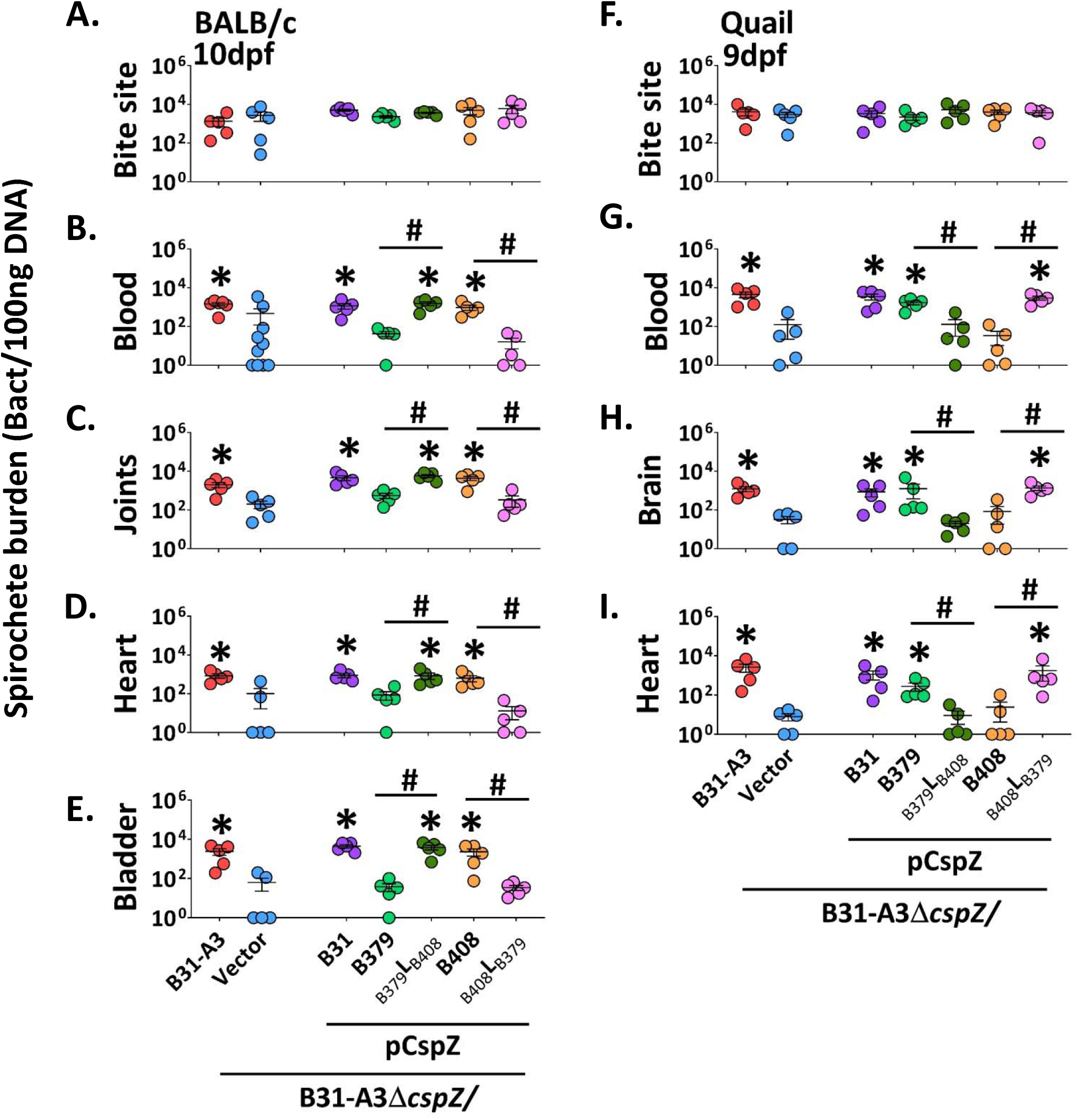
The CspZ loop-mediated FH-binding activity defines host-specific early hematogenous dissemination in a complement-dependent manner. *I. scapularis* nymphs carrying B31-A3, B31-A3Δ*cspZ* harboring the empty vector pKFSS (“Δ*cspZ*/Vector”), or this *cspZ* mutant strain producing CspZ_B31_, CspZ_B379_, CspZ_B379_L_B408_, CspZ_B408_, or CspZ_B408_L_B379_ were allowed to feed until repletion on **(A to E)** BALB/c mice or **(F to I)** quail. The bacterial loads in the indicated distal tissues were determined by qPCR at 10 days post nymphs feeding (dpf) in mice and 9dpf in quail. The bacterial loads were normalized to 100ng total DNA. Shown are the geometric mean of bacterial loads ± SEM of five mice or quail per group, except for the blood from BALB/c mice, which has nine mice per group. Significant differences (p *<* 0.05, Kruskal-Wallis test with the two-stage step-up method of Benjamini, Krieger, and Yekutieli) in the spirochete burdens relative to the Δ*cspZ*/Vector (“*”) or between two strains relative to each other (“#”) are indicated.

### The population-wide CspZ loop structures evolved from a variant with versatile FH-binding ability

To investigate the evolutionary history of the CspZ loop structures, we mined *B. burgdorferi cspZ* from NCBI GenBank and sequence read archive (totaling 174 high-quality *cspZ* isolates originated from ticks, reservoir hosts, and patients, across North America, Europe, and Asia). Phylogenetic analysis revealed three lineages with uncertain grouping among them, suggesting insufficient variation of *cspZ* to allow for well-established phylogenetic groups to emerge (data not shown). However, a haplotype-based phylogenetic network distinctly separated these three individual lineages, each of which contains one of the above-tested *cspZ* alleles: *cspZ_B31_*, *cspZ_B379_*, or *cspZ_B408_* (**Fig. S9**). Isolates within the same lineage had over 99% sequence identity and the same loop structure compared to *cspZ_B408,_ cspZ_B379_*, or *cspZ_B31_* (containing the duplication, insertion, or neither at the loci encoding loop structure, respectively) (**Fig. 5A**). We pinpointed single amino acid polymorphisms (SAPs) in the isolates and found none of the SAPs within each lineage were located in the FH-binding interface (**Table S1**), with the exception of a SAP in one isolate in the CspZ_B379_ lineage (**Table S4**)). Rather, the majority of the SAPs were between lineages, and were located in the loop structures that are driving the FH-binding abilities. While this present study has been the only one to investigate the abilities of CspZ variants to bind house mouse and quail FH, the human FH-binding abilities of 13% of these 174 isolates have been evaluated (27, 28, 32, 41). Incorporating these findings with the phylogeny, we found a 100% correlation of known host-specific FH-binding ability with lineage (**Fig 5A**). These results suggest each CspZ lineage with distinct loop structures can be linked to CspZ-specific, host-dependent FH-binding ability.

**Figure 5.**
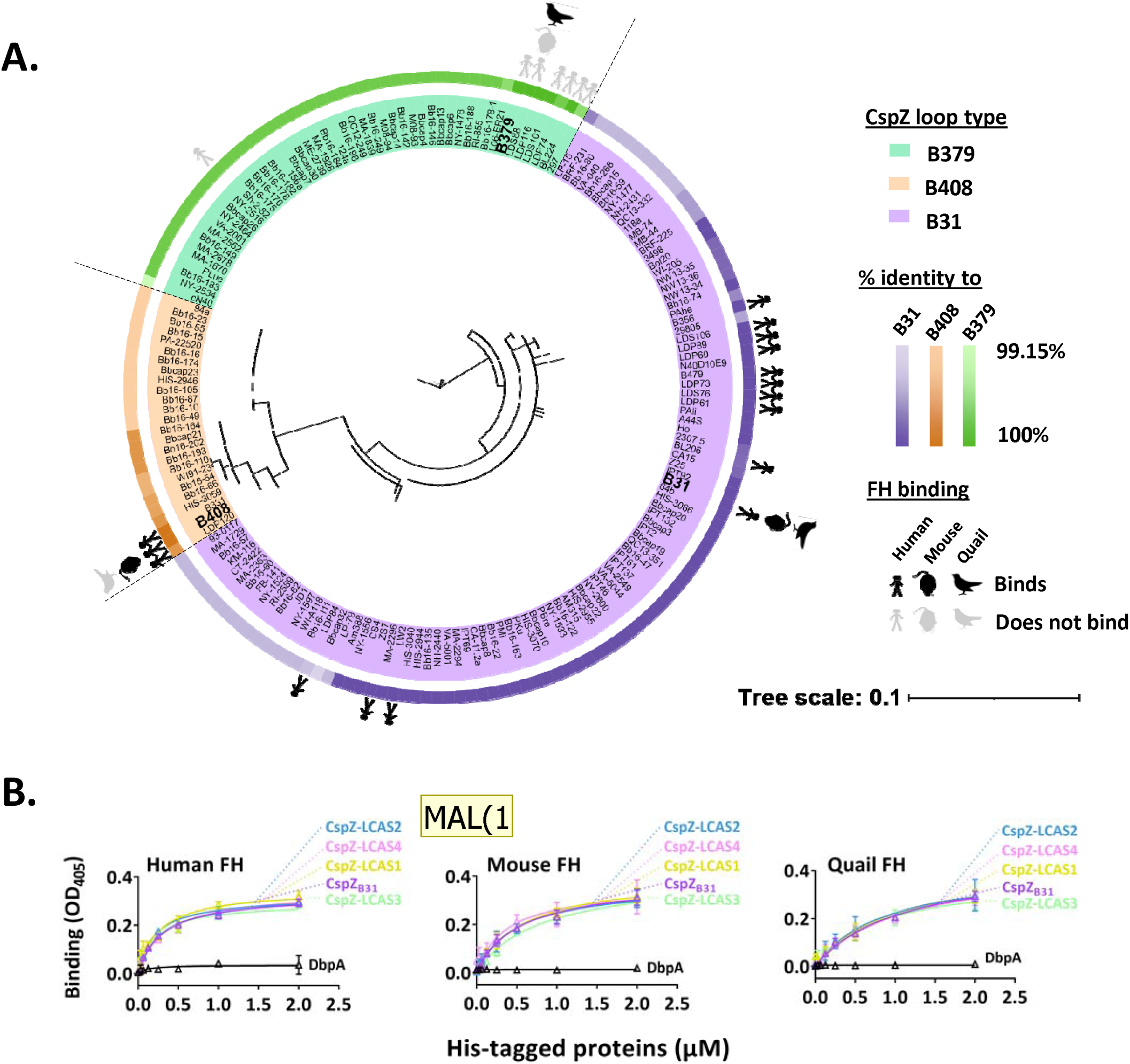
Phylogeny and sequence comparisons support polymorphic CspZ loop arising from host-specific adaptation of Lyme borreliae. (**A)** Unrooted likelihood tree of 174 *B. burgdorferi cspZ* isolates generated with IQ-tree and visualized in iTOL. The names of the isolates are highlighted based on their CspZ loop type (CspZ_B379_, CspZ_B408_, and CspZ_B31_ as green, orange, and purple, respectively). The percent identity of each isolate relative to CspZ_B31_ (purple), CspZ_B408_ (orange), or CspZ_B379_ (green) is indicated on the outer ring. The known FH-binding activity is marked: black or gray stick figures, mice, and quail indicate FH binding or lack of thereof, respectively, as determined herein and from previously published studies (27-29, 118, 119). **(B)** The indicated concentrations of recombinant histidine-tagged, predicted last common ancestor states of CspZ (“CspZ-LCAS1, CspZ-LCAS2, CspZ-LCAS3, and CspZ-LCAS4), CspZ_B31_, or DbpA (negative control) were added to triplicate wells coated with FH from human, mouse or quail, and protein binding was quantitated by ELISA. The experiments were performed with a single preparation of recombinant proteins tested in three independent iterations, in which samples were ran in duplicate. The K_D_ values **(Table S6)** representing the FH-binding affinity of each protein were determined from the average of three experiments. Shown is a representative iteration averaging the duplicates.

Additionally, we explored the evolutionary mechanisms that could have led to this minor loop structure-dependent FH-binding specificity. No evidence was found for recombination or past changes to the effective population size (data not shown). Although there was no evidence of selection for *cspZ* in its entirety (data not shown), several codons were undergoing either positive or negative selection (**Fig. S1A**). We reconstructed the possible last common ancestor states (“LCAS”) of the entire *B. burgdorferi* CspZ population using the chromosome mutation rate estimated in a previous study (42). We found that the diversification of the three lineages was estimated to be 261 to 784 years ago (y.a.) whereas the diversification of the LCAS likely occurred approximately 2166y.a. (896-5718 HPD_5-95%_) (**Table S5)**. The LCAS variants have a loop region resembling that of CspZ_B31_ both in AAs and structure (**Fig. S1A, S10**), and versatilely bound to human, mouse, and quail FH, similar to CspZ_B31_ (**Fig. 5B, Table S6**). These results suggest the diversification of CspZ is arisen relatively recently from a generalist variant with versatile FH-binding features.

## DISCUSSION

Host tropism is an outcome of ongoing host-pathogen interactions (15). In vector-borne zoonotic pathogens, such interactions include vector-to-host transmission, pathogen dissemination and persistence, and host-to-vector acquisition (43). For Lyme borreliae to survive throughout these infection steps, spirochetes need to overcome numerous host immunological mechanisms, including complement-mediated killing (17, 18, 44). In fact, Lyme borreliae inactivate complement in the tick blood meal for tick-to-host transmission and in the host bloodstream for dissemination and persistence (32, 45–47). In this study, we examined this inactivation conferred by a Lyme borreliae FH-binding protein, CspZ. We showed early colonization defects of a tick- introduced, *cspZ*-deficient mutant at distal tissues, defining this polymorphic protein as a contributor to the early stages of dissemination. Superimposing the crystal structures of three polymorphic CspZ variants from genotypically-diverse *B. burgdorferi* strains revealed a variable motif of a short AA stretch (30). Our newly-resolved and software-predicted complex structures of CspZ with human, mouse, and quail SCR6-7 further linked those loop-encoded variable residues to distinct host-specific FH binding activity. These results are congruent with the ability of CspZ variants and their associated mutants with swapped loop structures to promote host-specific FH-binding activity, complement evasion, and dissemination, defining this CspZ loop structure as a determinant of host tropism. Our findings thus demonstrate the concept that minor variation could functionally impact host-adapted phenotypes, potentially modulating host tropism (7–9). Our findings can also be attributed to a structurally unique FH-binding mechanism, as the FH-binding interface of CspZ is significantly different from that of the only other structurally-characterized SCR6-7-binding pathogen protein, *Neisseria meningitidis* Fhbp (48)(**Fig. S11**). Taken together, these results provide a platform of using structure-guided approaches in identifying pathogens’ host tropism determinants and their biological functions.

We also integrated an evolutionary approach to investigate the emergence of such host tropism determinants, an approach whose importance has been recently demonstrated to understand and track modern disease emergence (e.g., (49, 50)). Our findings demonstrate that the *B. burgdorferi* CspZ variants diversified from a last common ancestor that could bind human, mouse, and quail FH. This diversification occurred relatively recently (**Table S10**), compared to the previous finding showing that *B. burgdorferi* as a species evolved about 60,000 years BP, and the other FH-binding protein, CspA, diversified about 25,000 years y.a. (42, 45). The fact that CspZ has no orthologs in non-*Borrelia* organisms also supports the young age of this protein, suggesting CspZ may be undergoing weaker purifying and more variable selective pressure (30, 51). We did not detect selective pressure on the entirety of the *cspZ* locus, but rather only in a few individual codons, none of which were in the loop structures driving FH-binding phenotypes. The loop structures, which are the most variable part of these CspZ variants, contain indels, so the detection of gene-wide selective pressures was likely masked by the exclusion of gaps by the selection tests. In fact, other polymorphic anti-complement loci, particularly *ospC*, are undergoing balancing selection (52), (53). Both Multiple Niche Polymorphism (MNP) and Negative Frequency Dependent Selection (NFDS) have been proposed as mechanisms driving this selection. On the one hand, NFDS proposes that a high diversity of alleles emerge through the negative correlation between spirochete fitness and the frequency of any allele in the overall population (54). In Lyme borreliae, NFDS may explain the greater antibody-mediated clearance of spirochete strains harboring *ospC* alleles of higher frequency loci (55, 56). On the other hand, MNP proposes that the diversity of alleles/genotypes is maintained by fitness variation of these spirochetes across reservoir hosts (53, 57). Though it is possible that lack of selective pressure detection reflects the neutral selection experienced in *cspZ*, as a systemically redundant anti-complement protein (58), the possibility that *cspZ* is undergoing the same selective pressures as other functionally redundant proteins (e.g., *ospC*) cannot be excluded. In fact, balancing selection on one locus can increase diversity in another genetically-linked locus (59), and there is genetic linkage between CspZ types and *ospC* types (27, 28, 36). This may raise an intriguing possibility that the balancing selection on *ospC* may allow diversification through drift in other functionally redundant genes like *cspZ* without deleterious effect to the spirochetes, which warrants further investigations.

Some Lyme borreliae species or strains carry determinants that promoting host-specific phenotypes that differ from the known host range of these species or strains (36, 60–62). For example, the *B. burgdorferi* strain 297 is highly infectious in mice, but its CspZ loop-structures are identical to those in CspZ_B379_ from the strain B379 that is not mouse adapted (36, 62). This reflects the fact that most conclusions from this study were drawn by using spirochete strains with the same genetic background but producing the CspZ variants of interest, which may not account for polygenic contributions of other proteins in different strains (36). One possibility that addresses this discrepancy is that host-adapted phenotypes are conferred by not only CspZ but other anti-complement proteins, and the functional contribution from each of these proteins differ (and/or are host-dependent). In fact, the colonization defects of the *cspZ*-deficient *B. burgdorferi* were not found in the initial infection site at early infection stages, or in the distal tissues at later timepoints, consistent with the concurrent production of other FH-binding proteins (25, 45, 63–65). Additionally, non-FH-binding anti-complement proteins (66–68) and/or tick salivary proteins with anti-complement functions (69–71) could also drive complement-based phenotypes. Further, pathogen evasion to complement-independent mechanisms (e.g., antibodies), or even the host immune evasion-independent adaptive phenotypes (e.g., adhesion), can occur simultaneously with the anti-complement mechanisms (72), all contributing to the overall host-adapted phenotypes (6). All these confounding factors may complicate the delineation of the roles for CspZ at later stages of infection. Thus, our results do not rule out the possibility of CspZ-mediated complement evasion at distal tissues during post-dissemination timepoints, but rather highlight the importance of this protein immediately after spirochetes begin to disseminate. Despite such caveats, the strength of these isogenic strains needs to be emphasized in delineating the roles of a particular pathogen determinant from an enormous complexity of functionally redundant proteins (24, 45, 73, 74). Such potential polygenic or multi-factorial, and host-specific phenotypes should consider the complexity of numerous determinants across different mechanisms in defining the effect on host tropism.

In this study, we used Lyme borreliae as a model to apply structural, microbiological, and evolutionary approaches to identify a determinant of host-adapted phenotypes and the specific mechanism impacting host tropism. It should be noted that distinct *B. burgdorferi* infectivity have been observed for host species within the same taxa (e.g., house mice vs. the reservoir white-footed mice) (6, 75, 76). Thus, we do not intend to extend the host-specific phenotypes from these laboratory models to every individual reservoir animal in the same host taxa. Rather, our results establish a platform with more accessibility of tools to facilitate the identification of host tropism determinants and their underlying mechanisms. Such results build the foundation to further examine similar concepts in reservoir animals to recapitulate field findings in understanding the mechanisms of host tropism in a controllable laboratory setting. The information and the platform established in this multi-disciplinary study establishes greater insights of pathogen-host interactions, facilitating the understanding of host tropism as the cause of newly emergent infectious diseases.

## MATERIALS AND METHODS

### Ethics statement

All mouse and quail experiments were performed in strict accordance with all provisions of the Animal Welfare Act, the Guide for the Care and Use of Laboratory Animals, and the PHS Policy on Humane Care and Use of Laboratory Animals. The protocol was approved by the Institutional Animal Care and Use Committee (IACUC) of Wadsworth Center, New York State Department of Health (Protocol docket number 19-451). All efforts were made to minimize animal suffering.

### Mice, quail, ticks, bacterial strains, animal sera, OmCI, and FH

BALB/c and Swiss Webster mice were purchased from Taconic (Hudson, NY). C3^-/-^ mice in the BALB/c background were generated from the C3^-/-^(C57BL/6) from Jackson Laboratory (Bar Harbor, ME) as described (24). *Coturnix* quail were purchased from Cavendish Game Birds Farm (Springfield, VT). *Ixodes scapularis* tick larvae were purchased from National Tick Research and Education Center, Oklahoma State University (Stillwater, OK) or obtained from the CDC through BEI Resources (Manassas, VA).

The *Escherichia coli, Pichia pastoris* and *Borrelia* strains used in this study are described in **Table S7**. *E. coli* strains DH5α, BL21(DE3), and derivatives were grown in LB broth or agar, supplemented with kanamycin (50µg/ml), ampicillin (100µg/ml), or no antibiotics as appropriate. *P. pastoris* strain X-33 was grown on YPD plates supplemented with zeocin (800µg/ml) or BMGY medium supplemented with 1% methanol. All *B. burgdorferi* strains were grown in BSK-II completed medium supplemented with kanamycin (200µg/mL), streptomycin (50µg/mL), or no antibiotics (**Table S7**).

Mouse FH was purchased from MyBiosource (San Diego, CA). Quail FH and recombinant OmCI proteins were generated as described previously (24, 32, 36, 39). The mouse and quail sera were obtained from Southern Biotech, Inc (Birmingham, AL) and Canola Live Poultry Market (Brooklyn, NY), respectively. The sera from white-footed mice were obtained previously (6). Prior to being used, all these sera were screened for antibodies against the C6 peptide of the *B. burgdorferi* protein VlsE (77) with the C6 Lyme ELISA kit (Diamedix, Miami Lakes, FL) to ensure the mice did not have prior exposure to *B. burgdorferi*.

### Generation of recombinant CspZ proteins and recombinant human FH SCR6-7

To generate recombinant CspZ proteins for crystallization, *cspZ_B379_* (GenBank: FJ911671.1) and *cspZ_B408_* (GenBank: FJ911677.1) were amplified by PCR from the genomic DNA of *B. burgdorferi* strain B379 and B408 using the primers listed in **Table S8**. Note that B408 has two copies of *cspZ*, but they are functionally identical with only a single synonymous SNP at nucleotide 699 (36, 78). Based on the prediction by SignalP 4.1 (79) and according to our previous structural data from CspZ_B31_ (30), the lipoprotein signal peptide (residues 1-22) was excluded from the amplified gene. The introduced *NcoI* and *NotI* restriction sites were used for ligation of the amplified fragments into the pETm-11 expression vector which contains the coding region for an N-terminal 6xHis tag and a tobacco etch virus (TEV) protease cleavage site. Expression in *E. coli*, purification by affinity chromatography, and 6xHis tag cleavage by TEV protease of both proteins CspZ_B379_ and CspZ_B408_ were performed similarly as described previously for CspZ_B31_ (30). The purified and cleaved proteins were buffer exchanged into 10 mM Tris-HCl (pH 8.0) and concentrated to 11 mg/ml using an Amicon centrifugal filter unit (Millipore, Burlington, MA, USA).

To produce recombinant human FH for crystallization, the gene encoding the SCR6-7 of human FH was synthesized by BioCat GmbH (Heidelberg, Germany) and cloned into pPICZαA vector behind the α-factor secretion signal using *XhoI* and *NotI* restriction sites in a way to restore the Kex2 signal cleavage site. The plasmid was linearized with *PmeI* and transformed by electroporation into *Pichia pastoris* (reassigned as *Komagataella phaffii*) strain X-33. Transformants were obtained on YPD agar plates containing 800µg/ml of the antibiotic zeocin. The selected clone was cultivated 24-h in BMGY medium at 30°C with aeration (250rpm) following addition of 1% methanol daily, and cultivation was continued for three more days. The cell pellet was removed by low-speed centrifugation. Supernatant was buffer-exchanged into 50mM sodium phosphate (pH 6.0) by Sephadex G-25 Fine column (bed volume 360ml) (Cytiva, Marlborough, MA, USA) in 100ml portions at a flow rate of 20ml/min. Two liters of supernatant was passed through the CaptoS Improved Resolution column (bed volume 20ml) (Cytiva, Marlborough, MA, USA) and bound material was eluted with a linear salt gradient at a flow rate of 6ml/min. Target protein fractions were selected based on SDS-PAGE. The relevant fractions were pooled and buffer-exchanged into 20mM Tris-HCl (8.0), 50mM NaCl and 10mM NaH_2_PO_4_ using an Amicon filter device (Millipore, Burlington, MA, USA).

To generate recombinant CspZ_B379_ and CspZ_B408_ for the studies other than crystallization, the region encoding *cspZ_B379_* or *cspZ_B408_* without the signal peptide was amplified as described above and engineered to encode *BamHI* and *SalI* sites at the 5’ and 3’ ends, respectively, allowing subsequent cloning into the pJET cloning vector (Thermo Fisher Scientific, Waltham, MA). These pJET-derived plasmids encoding *cspZ_B379_* or *cspZ_B408_* were used as template for site-directed, ligase-independent mutagenesis (SLIM) (**Table S7**) to generate plasmids producing CspZ_B379_- L_B408_ and CspZ_B408_-L_B379_ (80). After verifying the sequences of all the plasmids (Wadsworth ATGC facility), the DNA fragments were subsequently excised using *BamHI* and *SalI* and then inserted into the same sites in pGEX4T2 (GE Healthcare, Piscataway, NJ) (32). The pGEX4T2- derived plasmids were then transformed into the *E. coli* strain BL21(DE3). The GST-tagged CspZ proteins were produced and purified by affinity chromatography. These proteins were verified for their secondary structures not impacted by the mutagenesis using CD (**Fig. S12**), as described in the section “Circular dichroism (CD) spectroscopy.”

To generate recombinant CspZ from the last common ancestor states, pET-28a+ encoding these states flanked by *BamHI* and *SalI* sites at the 5’ and 3’ ends, respectively, were cloned (Synbio Technologies, Monmouth Junction, NJ). The plasmids were transformed into the *E. coli* strain BL21(DE3), and the His-tagged CspZ proteins were produced and purified by affinity chromatography.

### Crystallization and structure determination

For crystallization of CspZ_B379_ and CspZ_B408_, 96- well sitting drop plates were set using a Tecan Freedom EVO100 workstation (Tecan Group, Männedorf, Switzerland) by mixing 0.4μl of protein with 0.4μl of precipitant using the 96-reagent sparse-matrix screens JCSG+ and Structure Screen 1&2 (Molecular Dimensions, Newmarket, UK). The crystals for CspZ_B379_ were obtained in 0.2M Ammonium citrate and 24% PEG 3350. For CspZ_B408_, the crystals were formed in 0.2M potassium acetate, 0.1M Tris-HCl (pH 8.0) and 28% PEG 3350. Prior to the data collection, the crystals were frozen in liquid nitrogen. An additional 20% glycerol was used as a cryoprotectant for CspZ_B379_ crystals, whereas the respective precipitant with an additional 14% glycerol was used as cryoprotectant for CspZ_B408_ crystals.

CspZ_B408_ (4mg/ml) and human SCR6-7 (3mg/ml) were mixed together at a molar ratio of 1:2 and loaded on a HiLoad 16/600 Superdex 200 prep grade column (GE Healthcare, Chicago, IL, USA) pre-equilibrated with 20mM Tris-HCl (8.0), 50mM NaCl and 10mM NaH_2_PO_4_. The flow rate was set to 2ml/min. Size exclusion chromatography resulted in one major peak containing the complex, confirmed by SDS-PAGE. Crystallization was set as described earlier for CspZ_B379_ and CspZ_B408_ by mixing 0.4μl of protein with 0.4μl of precipitant and using the 96-reagent sparse-matrix screens. The crystals for CspZ_B408_-SCR6-7 complex were obtained in 0.2M Zinc acetate, 0.1M imidazole (pH 7.4) and 10% PEG 3000. Crystals were frozen in liquid nitrogen by using 20% glycerol as a cryoprotectant.

Diffraction data for CspZ_B379_, CspZ_B408_ and CspZ_B408_-SCR6-7 complex were collected at the MX beamline instrument BL 14.1 at Helmholtz-Zentrum, Berlin (81). Reflections were indexed by XDS and scaled by AIMLESS from the CCP4 suite (82–84). Initial phases for CspZ_B379_ and CspZ_B408_ were obtained by molecular replacement using Phaser (85), with the crystal structure of the orthologous protein CspZ_B31_ was used as a searching model (97% sequence identity, PDB: 4CBE). For CspZ_B408_-SCR6-7 complex, the phases were determined using CspZ_B408_ (PDB: 7ZJK, RMSD 0.98 Å) and human FH SCR6-7 (PDB: 4AYD-A, RMSD 0.98 Å) as the searching models. After molecular replacement, the protein models were built automatically in BUCCANEER (86). The crystal structures were improved by manual rebuilding in COOT (87). Crystallographic refinement was performed using REFMAC5 (88). A summary of the data collection, refinement and validation statistics for CspZ_B379_, CspZ_B408_ and CspZ_B408_-SCR6-7 complex are given in **Table S9.**

### Protein 3D structure prediction using AlphaFold

AlphaFold v2.0 (34) was used to predict the 3D structure for quail FH SCR6-7 extrapolated from the sequences of *Coturnix japonica* complement FH (GenBank: XM_015869474.2). Structure prediction with AlphaFold v2.0 was performed according to the default parameters as indicated at the website https://github.com/deepmind/alphafold/ running on AMD Ryzen Threadripper 2990WX 32-Core; 128 GB RAM; 4 x NVIDIA GeForce RTX 2080, and using the full databases downloaded on 2021-09-25. For further structural analysis, only the predicted structure with the highest confidence was used (as ranked by using LDDT (pLDDT) scores).

### Circular dichroism (CD) spectroscopy

CD analysis was performed on a Jasco 810 spectropolarimeter (Jasco Analytical Instrument, Easton, MD) under nitrogen. CD spectra were measured at room temperature (RT, 25°C) in a 1mm path length quartz cell. Spectra of each of the CspZ proteins (10μM) were recorded in phosphate based saline buffer (PBS) at RT, and three far-UV CD spectra were recorded from 190 to 250nm for far-UV CD in 1nm increments. The background spectrum of PBS without proteins was subtracted from the protein spectra. CD spectra were initially analyzed by the software Spectra Manager Program (Jasco). Analysis of spectra to extrapolate secondary structures were performed using the K2D3 analysis programs (89).

### ELISAs

Quantitative ELISA was used to determine FH-binding by CspZ proteins, or ancestral proteins, as described previously (24, 90), with the following modifications: Mouse anti-GST tag or mouse anti-His tag 1:200× (Sigma-Aldrich) and HRP-conjugated goat anti-mouse IgG 1:2,000× (Seracare Life Sciences) were used as primary and secondary antibodies, respectively, to detect the binding of GST- or histidine-tagged proteins.

### Surface Plasmon Resonance (SPR)

Interactions of CspZ proteins with FH were analyzed by SPR using a Biacore T200 (Cytiva, Marlborough, MA). Ten micrograms of mouse or quail FH were conjugated to a CM5 chip (Cytiva) as described previously (90). For quantitative SPR experiments, 10µL of increasing concentrations (0.08, 0.03125, 0.0125, 0.5, 2µM) of each of the CspZ proteins were injected into the control cell and the flow cell immobilized with FH at 10μl/min, 25°C. To obtain the kinetic parameters of the interaction, sensogram data were fitted by means of BIAevaluation software version 3.0 (GE Healthcare), using the one step biomolecular association reaction model (1:1 Langmuir model), resulting in optimum mathematical fit with the lowest Chi-square values.

### Shuttle vector construction and plasmid transformation into *B. burgdorferi*

“Loop swapped” CspZ variants (i.e., CspZ_B379_L_B408_ and CspZ_B408_L_B379_) were designed based on the full-length sequences (B379 accession: OM643341; B408: accession: OM643340) and purchased as double-stranded DNA fragments flanked by *BamHI* and *SalI* on the 5’ and 3’, respectively (Integrated DNA Technologies, Inc., Coralville, IA). B31-A3Δ*cspZ* was complemented with these variants, or with native CspZ from B379 and B408 flanked by the same restriction enzyme sites (**Table S7**), in the same manner as the previously published strains of B31-A3Δ*cspZ*/pKFSS and B31- A3Δ*cspZ*/pCspZ_B31_ (32). The plasmid profiles of these spirochetes were examined to ascertain identical profiles between these strains and their parental strain B31-A3 (91). The generation time of these transformants was calculated as previously described (32).

### Flow cytometry

CspZ production and FH-binding on spirochete surface were determined as described (24), including blood-treatment to induce the production of CspZ (32). To determine the levels of mouse C5b-9 or quail C8 deposition on the surface of spirochetes, mouse or quail sera were incubated with 1x10^7^ spirochetes in PBS at a final concentration of 20% at 25°C for one hour and the detection procedure has been described previously (36). Basically, after incubation, spirochetes were washed then resuspended in HBSC-DB (25mM Hepes acid, 150mM sodium chloride, 1mM MnCl_2_, 1mM MgCl_2_, 0.25mM CaCl_2_, 0.1% glucose, and 0.2% BSA). Rabbit anti-mouse C5b-9 polyclonal IgG (1:250x) (Complement Technology, Tyler, TX) or mouse anti-quail C8 polyclonal sera (1:250x) (36) were used as the primary antibodies. An Alexa 647-conjugated goat anti-rabbit (ThermoFisher) or a goat anti-mouse IgG (ThermoFisher) (1:250x) was used as the secondary antibody. After staining, the spirochetes were fixed with 0.1% formalin. The resulting fluorescence intensity of spirochetes was measured and analyzed by flow cytometry using a FACSCalibur (BD Bioscience) as described (32, 36).

### Serum resistance assays

The serum resistance of *B. burgdorferi* was measured as described with modifications (6, 24, 32). Cultures in mid-log phase of each strain treated with human blood (32), as well as the high passaged, non-infectious, and serum-sensitive human blood-treated *B. burgdorferi* strain B313 (control), were cultivated in triplicate and diluted to a final concentration of 5×10^6^ cells/mL in BSK-II medium without rabbit sera. The cell suspensions were mixed with sera collected from naïve white-footed mice or quail (60% spirochetes and 40% sera) in the presence or absence of 2µM of CVF (Complement Technology) or recombinant OmCI, to deplete complement from mouse and quail sera, respectively. Heat-inactivated sera (65°C for 2-h) were also included in each of the aforementioned combinations as a control (all strains survived equally; data not shown). To determine the bacteria survival, the number of motile spirochetes was counted under dark field microscopy at 0- and 4-h following incubation with sera as described (32), as we have shown that motile spirochetes determined using this methodology accurately reflect results using live-dead staining assays (39). The percent survival of *B. burgdorferi* was calculated by the normalization of motile spirochetes at 4-h post incubation to that immediately after incubation with sera.

### Mouse and quail infection by ticks

Generating flat, infected *I. scapularis* nymphs has been described previously (24, 92). The infected nymphs were placed in a chamber to feed on 4- to 6- week old male and female BALB/c or C3^-/-^ mice in BALB/c background, or on four- to six-week old male and female untreated or OmCI-treated quail, as described previously (45). For OmCI-treatment, the quail were subcutaneously injected with OmCI (1mg/kg of quail) one day prior to the nymph feeding. The engorged nymphs were obtained from the chambers at four days post tick feeding. Animals were sacrificed and tissues collected from the mice at 7-, 10-, or 14- (blood, tick bite site of the skin, heart, bladder, tibiotarsus joint), and quail (blood, tick bite site of the skin, heart, brain) at 9-, or 14-, days post nymph feeding. To ensure OmCI was still functional at these timepoints, quail were subcutaneously injected with OmCI (1mg/kg of quail) or PBS buffer (control), and the sera were collected at 10 days post injection (equivalent to 11 days post nymph feeding). The lack of the ability of this serum to kill the sera-sensitive *B. burgdorferi* strain B313 (i.e., to ensure complement was still depleted) was assessed as described in the section “Serum resistance assays.” (**Fig. S13**)

### Quantification of spirochete burden

The DNA from tissues, blood, and ticks was extracted as described previously (45). qPCR was then performed to quantitate bacterial loads. Spirochete genomic equivalents were calculated using an ABI 7500 Real-Time PCR System (ThermoFisher Scientific) in conjunction with PowerUp SYBR Green Master Mix (ThermoFisher Scientific) based on amplification of the Lyme borreliae *recA* gene using primers BBRecAfp and BBRecArp (**Table S8**) with the amplification cycle as described previously (32). The number of *recA* copies was calculated by establishing a threshold cycle (Cq) standard curve of a known number of *recA* gene extracted from *B. burgdorferi* strain B31-A3, then comparing the Cq values of the experimental samples.

### Genomic analyses

To generate the *cspZ* phylogenetic trees, we mined all publicly available *cspZ* sequences on NCBI as of September 2021, including assembled genomes, nucleotides, and unassembled genomes on the SRA. To pull *cspZ* from unassembled genomes, the short reads were aligned to *cspZ* from B31, B379, or B408 with UGENE v39.0 using BWA-MEM at defaults (93, 94). Strains were removed from the analyses if the coverage was too low, there was evidence of PCR/sequencing errors (e.g., non-conserved homopolymer length) or evidence of multiple CspZ variants within one strain. All resulting 174 *cspZ* sequences, plus the outgroup strains (*B. spielmanii* A14s accession: EU272854.1; *B. afzelii* FEM4 accession: OM243915; *B. afzelii* VS461 accession: MN809989.1; *B. garinii* PBr accession: CP001307.1; *B. bissettii* DN127 accession: NC_015916.1; *B. bissettii* CO275 accession: JNBW01000464.1), were aligned by codons using TranslatorX (95). All isolates were collapsed into haplotypes in FaBox v1.61, and these haplotypes were used with the *B. bissettii* outgroup to generate a NeighborNet network in SplitsTree v4 (96, 97). Phylogenetic trees were estimated using likelihood as optimality criterion in IQ-tree v1.6.12 (98) and a full substitution model search procedure in ModelFinder (99). Internode branch support was estimated with 10,000 replicatess of both ultrafast bootstrapping and the SH-aLRT branch test (98–100). All resulting phylogenetic trees were visualized in iTOL v6.4.3 (101). The pairwise sequence similarity for each of the 174 *B. burgdorferi cspZ* isolates relative to CspZ_B31_, CspZ_B408_, or CspZ_B379_ was determined in MEGA-X with default settings (102). Putative recombination breakpoints were analyzed with GARD (103), and evidence of selection for individual codons or branches on the *cspZ* gene tree were inferred with FUBAR (104), FEL (105) and MEME (106), all on the Datamonkey server (107). Selection tests for the entire *cspZ* gene included Tajima’s neutrality test in MEGA-X (102) and all tests available in DnaSP v5 with 10,000 coalescent simulations (108), all at default settings. The ancestor state for the entire *B. burgdorferi* ingroup was reconstructed using the LG model in GRASP 2020.05.05 (109, 110), as well as FireProt-ASR with default settings (111) using both full and haplotype phylogenies, a multitaxon outgroup and solely *B. bissettii* as outgroup. Divergence dating was carried out in BEAST v1.10.4 (112) using the HKY+Γ_4_ substitution model(113, 114), a coalescent Bayesian skyline coalescent model, and a strict clock with a uniform prior on the substitution rate using a previously determined rate of 4.75e-06 substitutions/site/year(42). A Markov chain Monte Carlo chain length of 100 million steps was used with a 10,000-step thinning, resulting in effective sample sizes greater than 200, an indication of an adequate chain mixing. The analyses were ran in triplicate and combined after removing a 10% burn-in.

The amino acids encoding SCRs 6-7 from human (GenBank accession U56979.1), mouse (NM_009888.3), or quail (XM_015869474.2) FH or CspZ_B31_, CspZ_B379_, and CspZ_B408_, the loop-swapped variants, and the reconstructed ancestral CspZ sequences were aligned in MEGA-X using ClustalW with default settings, analyzed with ESPript v3.0, and visualized with Jalview v2.11.0 (102, 115, 116).

### Statistical analysis

Samples were compared using the Mann-Whitney *U* test or the Kruskal-Wallis test with the two-stage step-up method of Benjamini, Krieger, and Yekutieli (117).

### Accession numbers

The coordinates and the structure factors for CspZ_B379_, CspZ_B408_, and human SCR-CspZ_B408_ have been deposited in the Protein Data Bank with accession codes 7ZJJ, 7ZJK, and 7ZJM, respectively.

## ACKNOWLEDGEMENTS

We thank Simon Starkey, Quinton Smith, Deirdre Torrisi, and Joey Anderson from the Wadsworth Veterinary Sciences facility for animal husbandry; Richard Marconi, Utpal Pal, Patricia Rosa, Ira Schwartz, and Gary Wormser for providing the *B. burgdorferi* strains; David Vance and Nicholas Mantis for sharing the *E. coli* strain; Timothy Czajka for the assistance of SPR analysis; Nikhat Parveen, Klemen Strle, and Grace Chen for insightful comments; and Laurel Lown for assistance with running PCRs. We also appreciate Leslie Eisele and Renjie Song of Wadsworth Biochemistry and Immunology Core for CD spectroscopy, SPR, and flow cytometry; and Karen Chave of the Wadsworth Protein Expression Core for the help of the materials to purify factor H. We thank Wadsworth ATGC core for plasmid sequencing. Diffraction data for *B. burgdorferi* CspZ proteins and CspZ-human FH were collected on BL14.1 at the BESSY II electron storage ring operated by the Helmholtz-Zentrum, Berlin. We would particularly like to acknowledge the help and support of Manfred S. Weiss and Jan Wollenhaupt during the diffraction data collection. This work was supported by NSF IOS1754995 (SOK), NSF IOS1755286 (YPL, ALM, TAN, ADII, JS, and AC), DoD TB170111, NIHR21AI144891, NIH R21AI146381, New York State Department of Health Wadsworth Center Start-Up Grant (ALM, TAN, YPL), and LOEWE Center DRUID Novel Drug Targets against Poverty-Related and Neglected Tropical Infectious Diseases, project C3 (PK). The funders had no role in study design, data collection and analysis, decision to publish, or preparation of the manuscript. The authors have declared that no competing interests exist.

## Supplemental Figures

**Figure S1.**
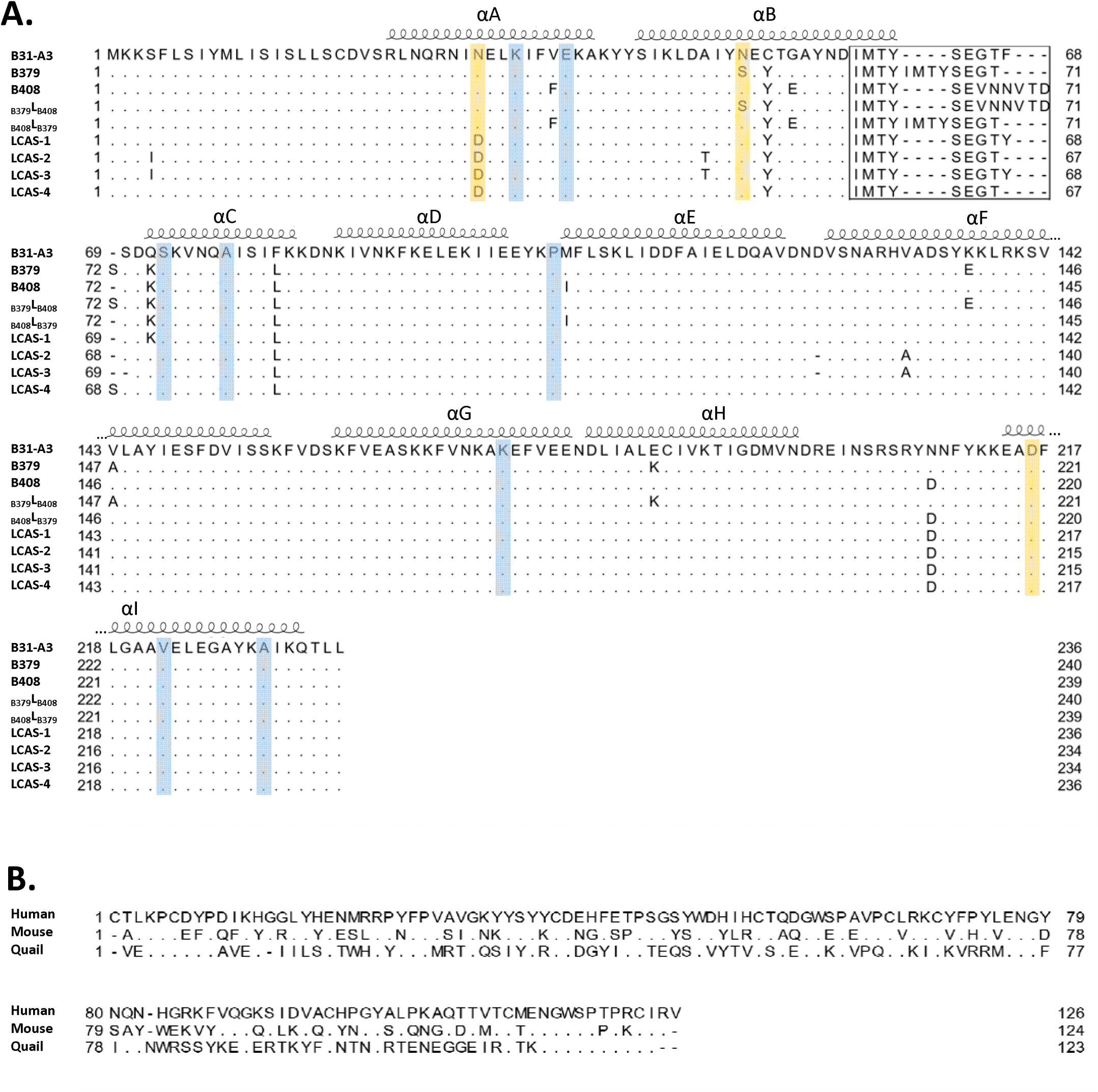
Amino acid alignments of the CspZ and FH variants. Amino acid alignments of **(A)** the indicated CspZ variants, mutants, and reconstructed ancestral states of variants or **(B)** human, mammalian, and avian FH SCR 6-7. (**A**) The CspZ amino acids accounting for the loop structures are indicated with the box, and the alpha-helices labeled above the sequences are extrapolated from the high-resolution structure of CspZ_B408_. The yellow and blue shading are indicative of loci showing evidence of positive and negative selection, respectively.

**Figure S2.**
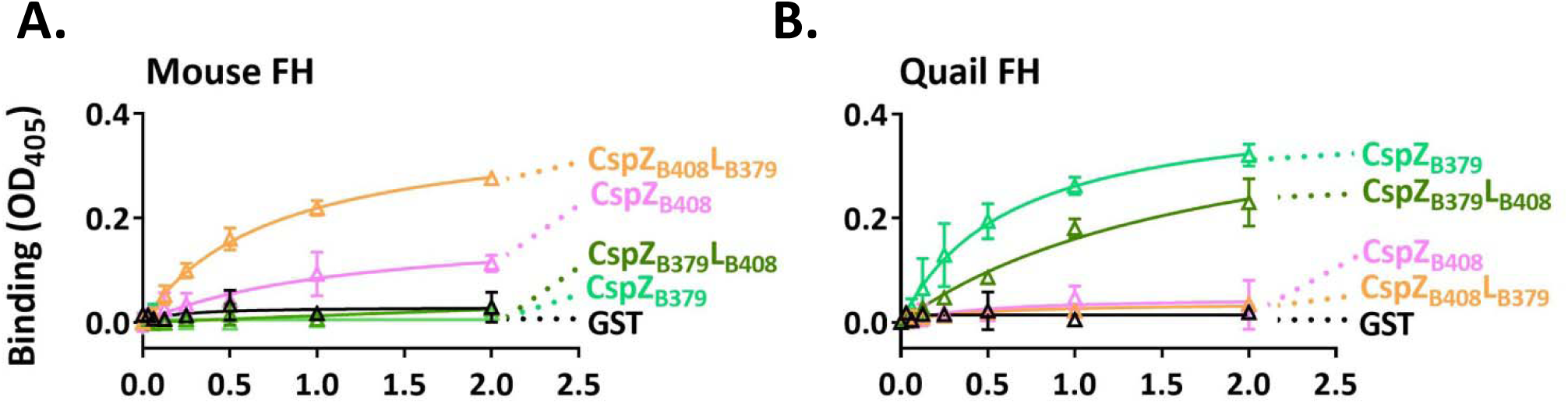
The polymorphic loop structures in recombinant CspZ proteins promote host-specific FH-binding ability determined by ELISA. The indicated concentrations of recombinant GST-tagged CspZ_B379_, CspZ_B408_, CspZ_B379_L_B408_, or CspZ_B408_L_B379_, or GST (negative control) were added to triplicate wells coated with FH from mouse or quail, and protein binding was quantitated by ELISA. The experiments were performed with a single preparation of recombinant proteins tested in three independent iterations, in which samples were ran in duplicate. The K_D_ values **(Table S2)** representing the FH-binding affinity of each protein were determined from the average of three experiments. Shown is a representative iteration averaging the duplicates.

**Figure S3.**
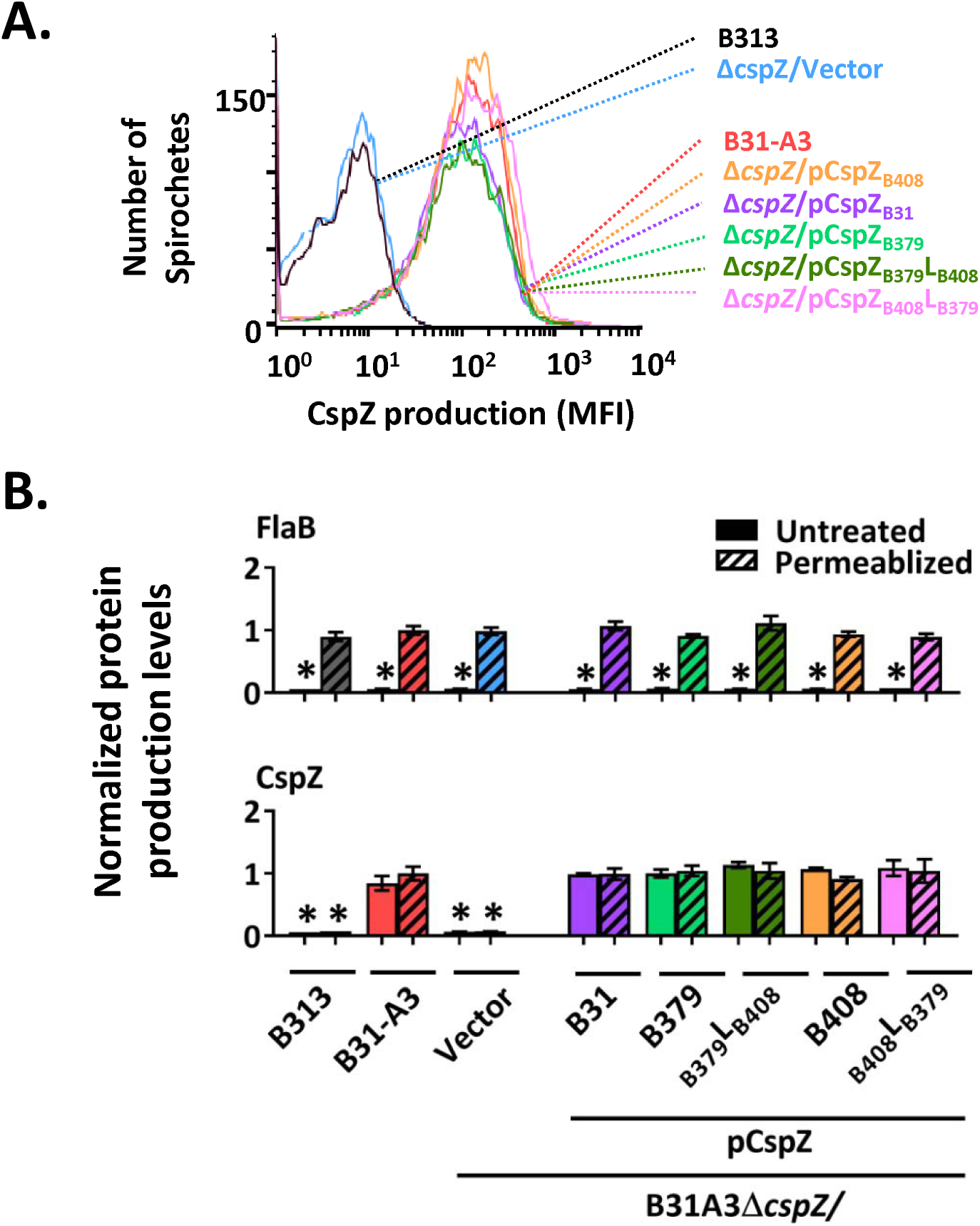
Indistinguishable surface production of CspZ among *B. burgdorferi* strains was observed using flow cytometry. Flow cytometry analysis of CspZ localized on the surface of *B. burgdorferi* strains B31-A3, B31-A3Δ*cspZ* harboring the vector pKFSS (“Δ*cspZ*/Vector”), or this *cspZ* mutant strain producing CspZ_B31_, CspZ_B379_, CspZ_B379_L_B408_, CspA_B408_, or CspZ_B408_L_B379_. **(A)** Representative histograms of flow cytometry analysis showing the levels of CspZ surface production on the indicated strains. **(B)** The production of FlaB (negative control) and CspZ on the surface of indicated *B. burgdorferi* strains was detected by flow cytometry. Values are shown normalized to the production levels of FlaB or CspZ on the surface of permeabilized B31-A3. Each bar represents the mean of four independent experiments ± the standard deviation. An asterisk (*) indicates that relative surface production of the indicated proteins was significantly lower (p *<* 0.05, Kruskal-Wallis test with the two-stage step-up method of Benjamini, Krieger, and Yekutieli) than that of permeabilized FlaB or CspZ by B31-A3.

**Figure S4.**
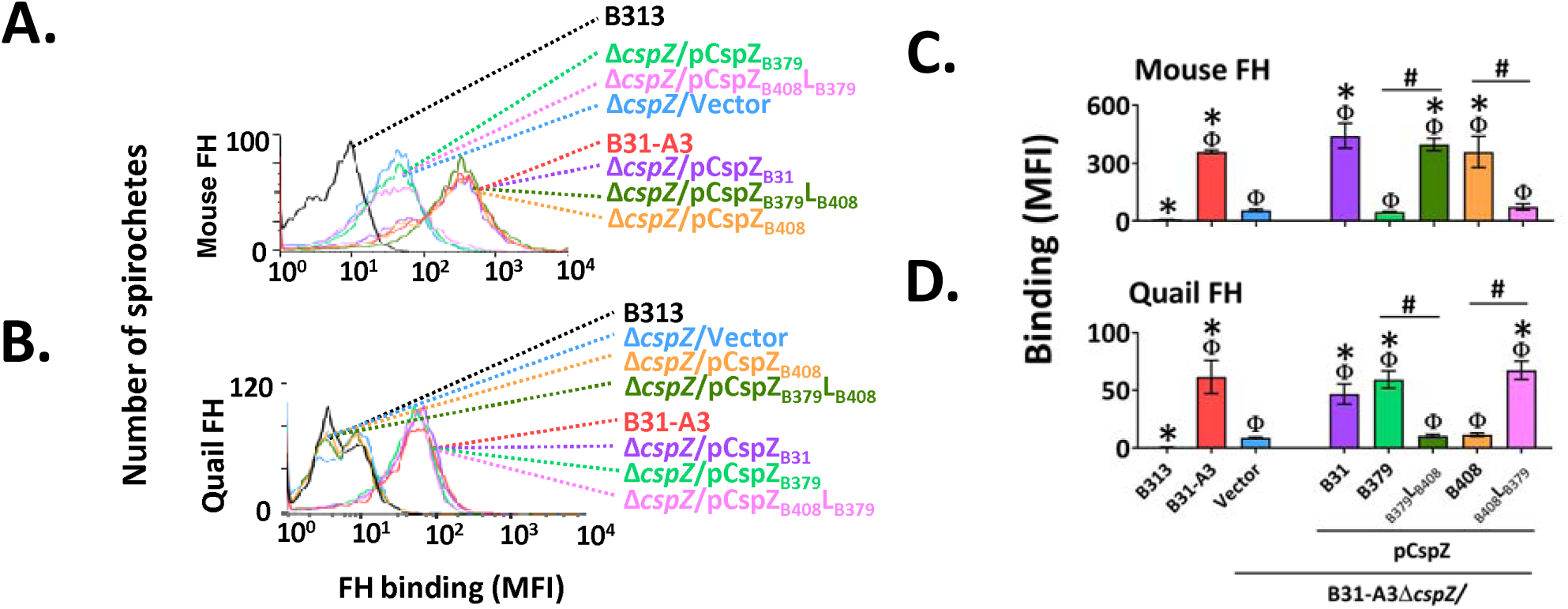
The polymorphic CspZ loop determines the host-specific, allelically variable FH-binding activity on spirochete surface. *B. burgdorferi* strains B313 (negative control), B31-A3, B31-A3Δ*cspZ* harboring the empty vector pKFSS (“Δ*cspZ*/Vector”), or this mutant strain producing CspZ_B31_, CspZ_B379_, CspZ_B379_L_B408_, CspZ_B408_, or CspZ_B408_L_B379_, was incubated with mouse or quail FH. The bacteria were stained with antibodies that recognize these FH variants prior to flow cytometry. Shown are the representative histograms of flow cytometry analysis presenting the levels of FH from **(A)** mouse or **(B)** quail binding to each *B. burgdorferi* strain. The levels of **(C)** mouse or **(D)** quail FH-binding were measured by flow cytometry and presented as mean fluorescence index (MFI). Each bar represents the mean of three independent experiments ± SEM. Significant differences (p *<* 0.05, Kruskal-Wallis test with the two-stage step-up method of Benjamini, Krieger, and Yekutieli) in the levels of FH-binding relative to the B313/Vector (“Φ”), Δ*cspZ*/Vector (“*”), or between two strains relative to each other (“#”) are indicated.

**Figure S5.**
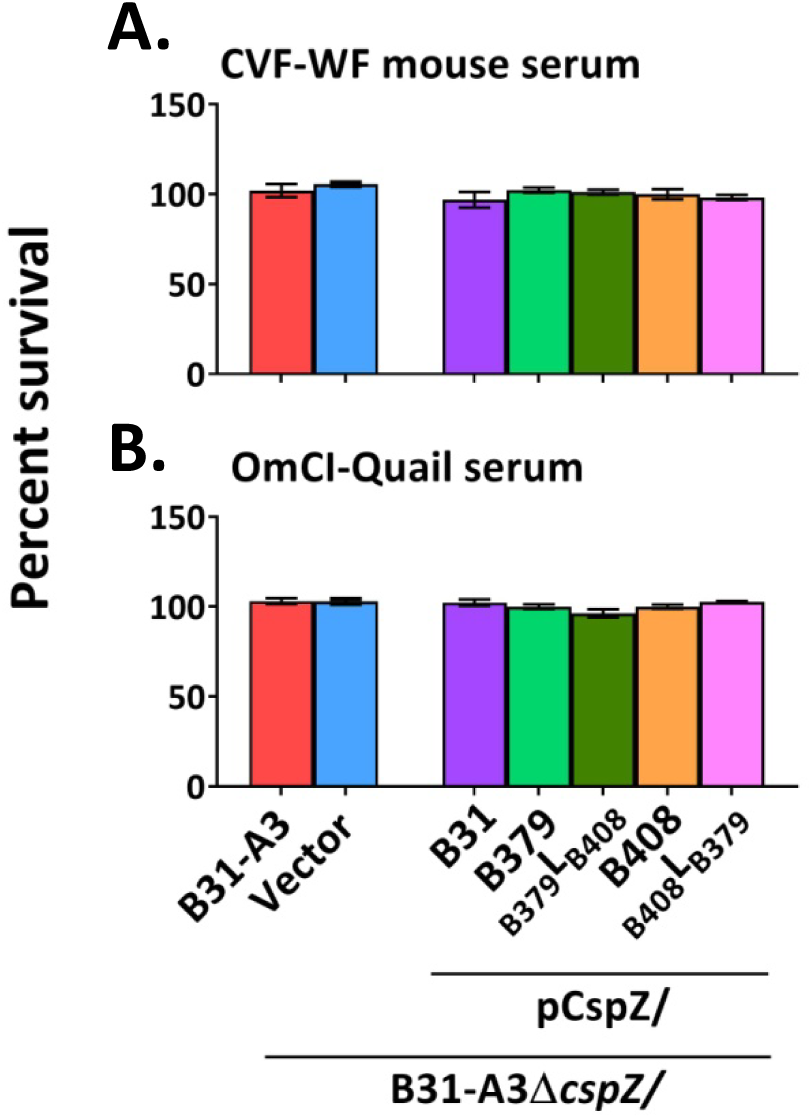
The CspZ loop-driven, host-specific serum resistance is recovered in complement-depleted sera. *B. burgdorferi* strains B313 (negative control), B31-A3, B31-A3Δ*cspZ* harboring the empty vector pKFSS (“Δ*cspZ*/Vector”), or this mutant strain producing CspZ_B31_, CspZ_B379_, CspZ_B379_L_B408_, CspZ_B408_, or CspZ_B408_L_B379_were incubated for 4-h with (**A)** CVF-treated white- footed mouse or **(B)** OMCI-treated quail sera, to a final concentration of 40%. The number of motile spirochetes was assessed microscopically. The precent survival of the *B. burgdorferi* strains was calculated using the number of motile spirochetes at 4-h post incubation normalized to that prior to incubation with sera. Each bar represents the mean of three independent experiments ± SEM. There were no significant differences (p *<* 0.05, Kruskal-Wallis test with the two-stage step-up method of Benjamini, Krieger, and Yekutieli) between the percent survival of any strain.

**Figure S6.**
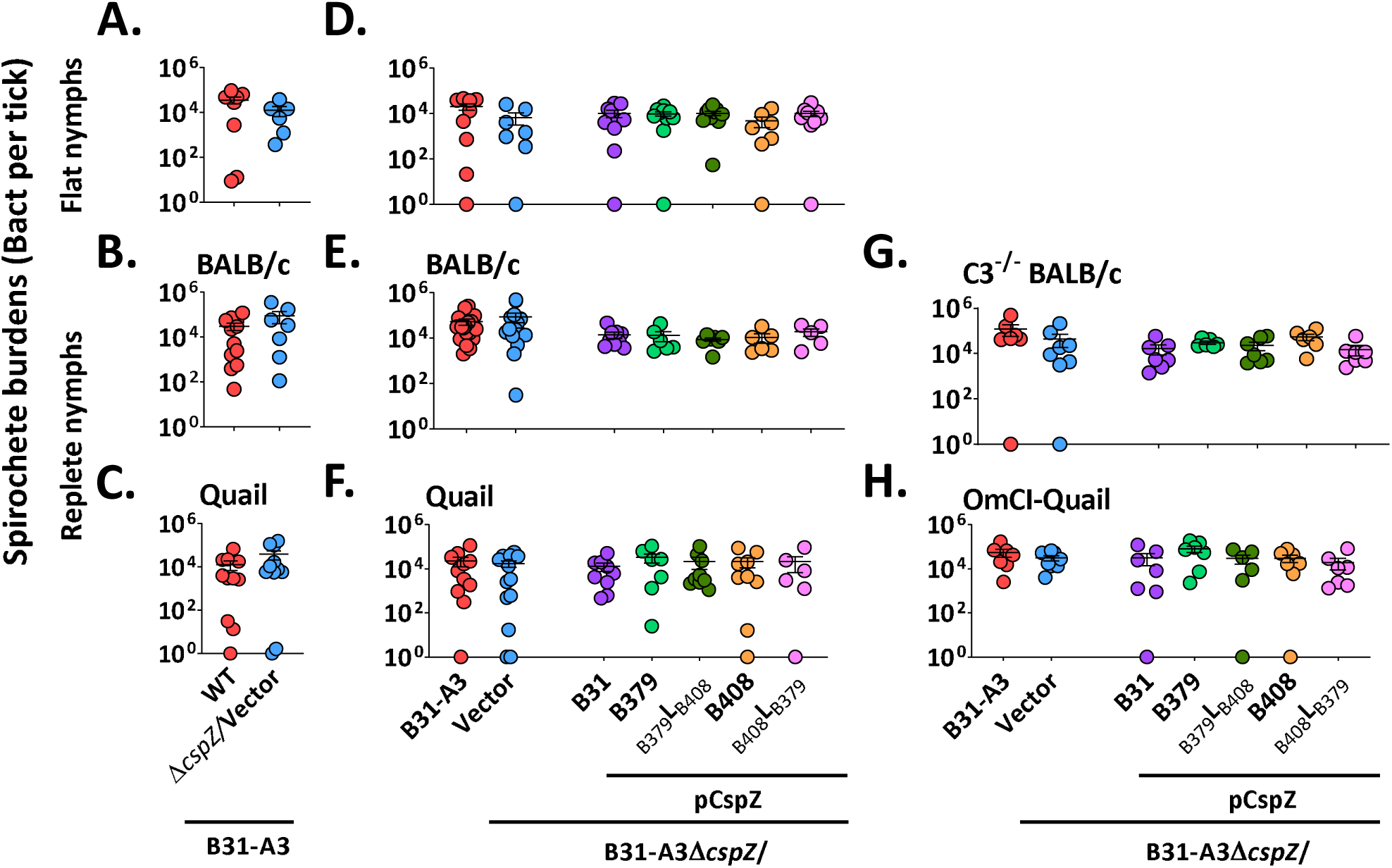
*B. burgdorferi* strains exhibit similar burdens in flat and fed nymphs. *B. burgdorferi*-infected flat nymphs were allowed to feed to repletion on **(B, E)** BALB/c or **(G)** C3-/- BALB/c mice, or **(C, F)** quail or **(H)** OmCI-treated quail. The spirochete loads in **(A, D)** flat or **(B, C, E, F, G, H)** replate nymphs were determined by qPCR. Shown are the geometric mean ± geometric standard deviation of at least five nymphs per group. There was no statistical difference (p > 0.05) of the spirochete burdens between different groups of the replete ticks using a **(A to C)** Mann-Whitney test or **(D to H)** Kruskal-Wallis test with the two-stage step-up method of Benjamini, Krieger, and Yekutieli.

**Figure S7.**
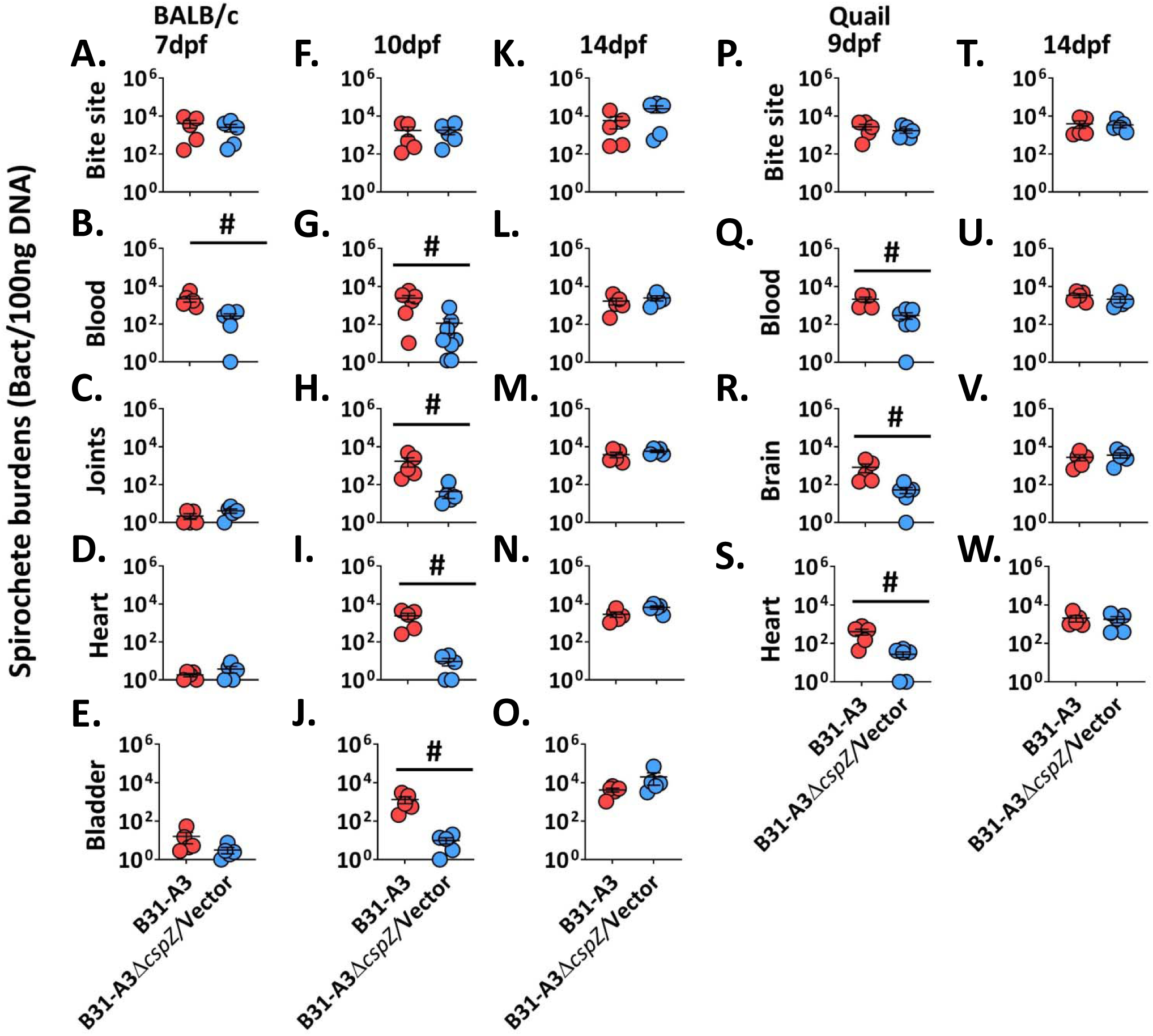
CspZ facilitates early bacteremia and distal tissue colonization during tick infection. The *I. scapularis* nymphs carrying *B. burgdorferi* strain B31-A3 or B31-A3Δ*cspZ* harboring the vector pKFSS (“Δ*cspZ*/Vector”) were allowed to feed until repletion on **(A to O)** BALB/c mice or **(P to W)** quail. The mice were euthanized at **(A to E)** 7, **(F to J)** 10, or **(K to O)** 14 days post nymphs feeding (dpf), whereas the quail were euthanized at **(P to S)** 9 or **(T to W)** 14dpf. **(A, F, K)** The site of the skin where nymphs fed (“Bite site”), **(B, G, L)** blood, **(C, H, M)** tibiotarsus joints, **(D, I, N)** heart, and **(E, J, O)** bladder of mice; and **(P, T)** the bite site, **(Q, U)** blood, **(R, V)** brain, and **(S, W)** heart of quail, were collected immediately after euthanasia and spirochete loads were determined by qPCR. The burdens were normalized to 100ng total DNA. Shown are the geometric mean of bacterial loads ± SEM of five mice or quail per group, except for samples from the mouse blood at 7 and 10dpf where the results are from six and nine mice, respectively. Significant differences (p < 0.05, Mann-Whitney test) in the spirochete burdens between two strains relative to each other (“#”) are indicated.

**Figure S8.**
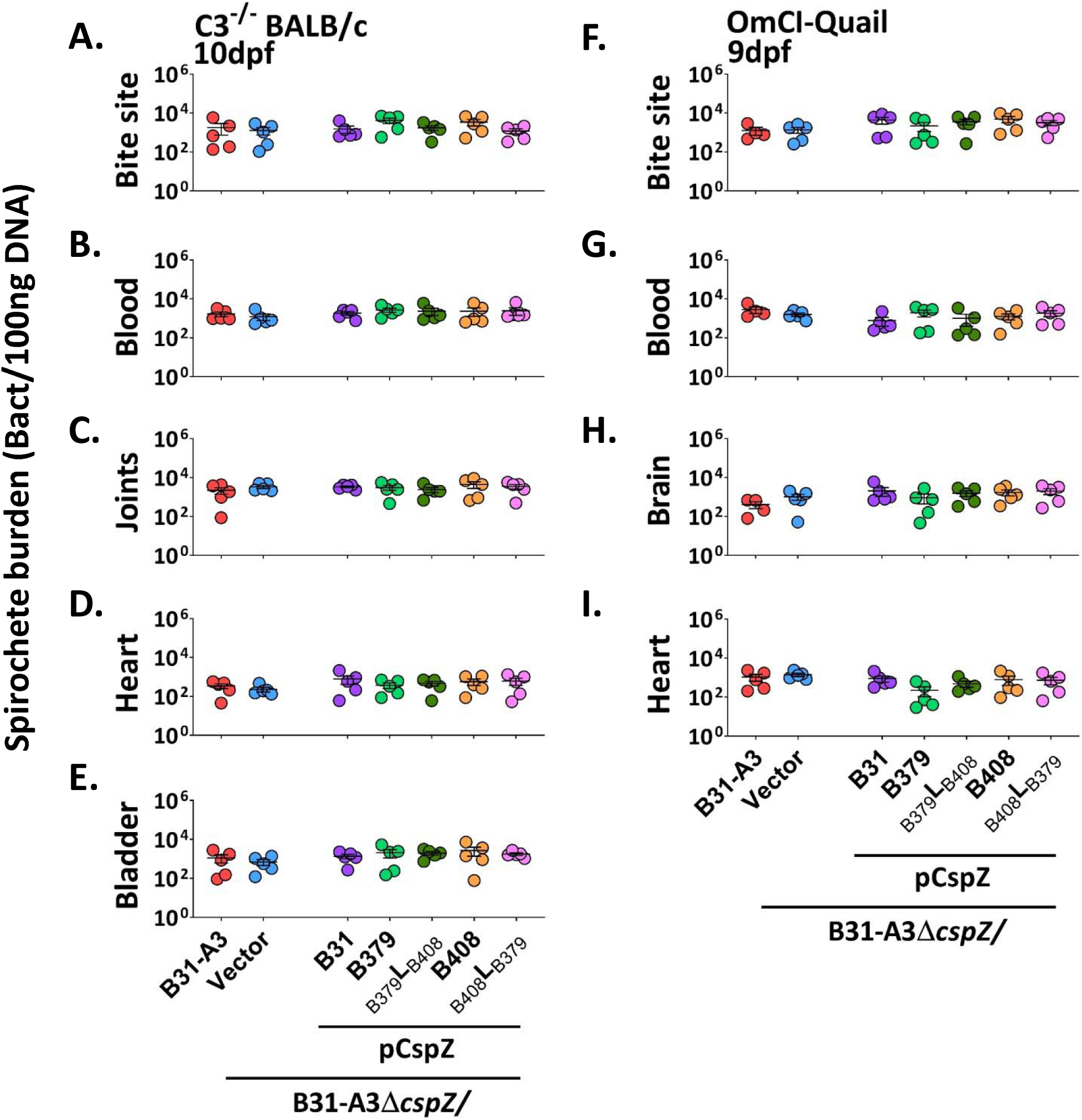
The CspZ loop-mediated early hematogenous dissemination is recovered when complement is depleted from hosts. *I. scapularis* nymphs carrying B31-A3, B31-A3Δ*cspZ* harboring the empty vector pKFSS (“Δ*cspZ*/Vector”), or this *cspZ* mutant strain producing CspZ_B31_, CspZ_B379_, CspZ_B379_L_B408_, CspZ_B408_, or CspZ_B408_L_B379_ were allowed to feed until repletion on **(A to E)** C3^-/-^ mice in a BALB/c background or **(F to I)** OMCI-treated quail, both of which deplete complement in the respective hosts. The bacterial loads in the indicated distal tissues were determined by qPCR and were normalized to 100ng total DNA, at 10 days post nymphs feeding (dpf) in mice and 9dpf in quail. Shown are the geometric mean of bacterial loads ± SEM of five mice or quail per group, except for the blood from BALB/c mice, which has nine mice per group. There were no significant differences (p *<* 0.05, Kruskal-Wallis test with the two-stage step-up method of Benjamini, Krieger, and Yekutieli) in the spirochete burdens for any strain.

**Figure S9.**
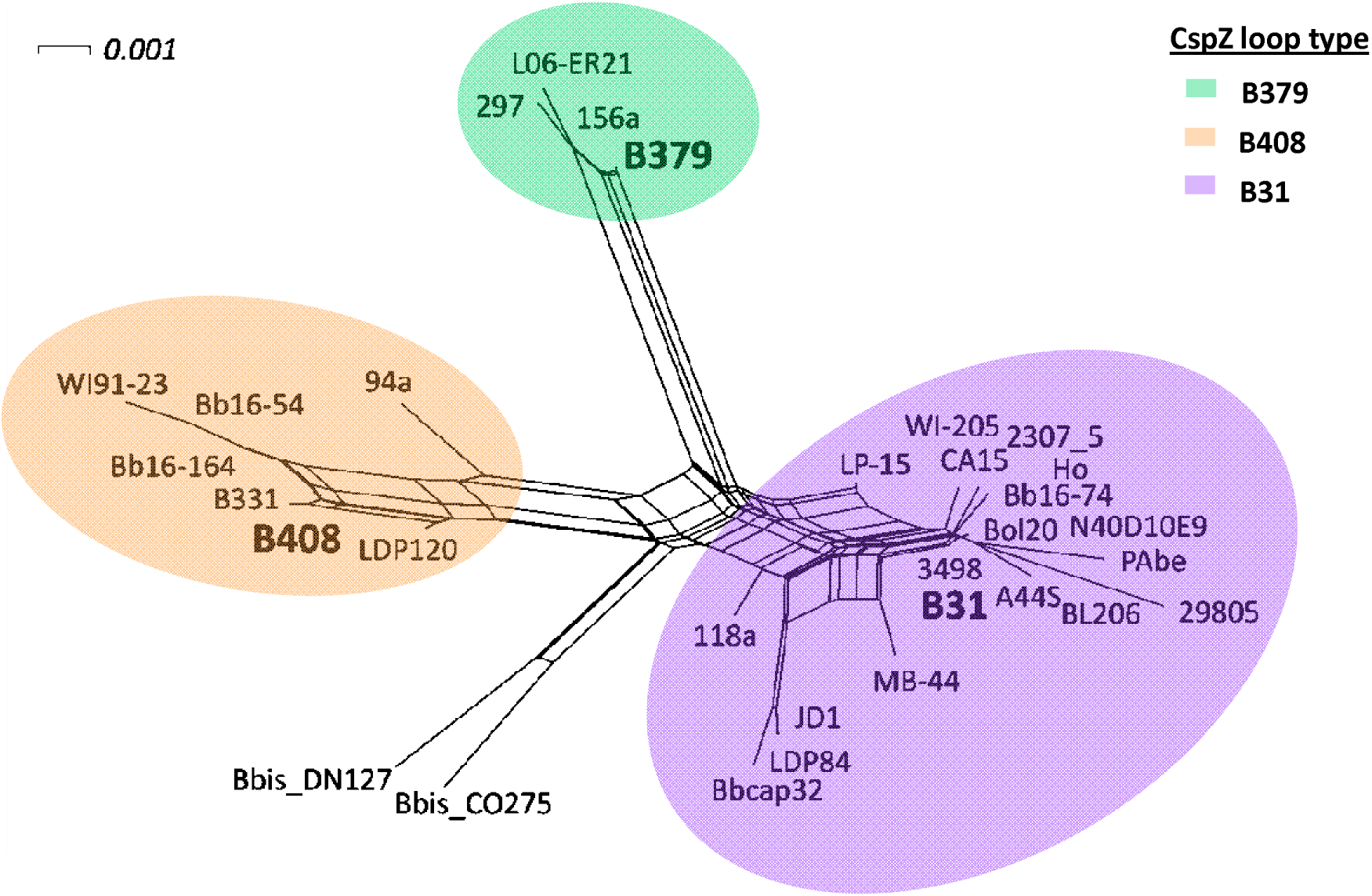
Phylogenetic network of *cspZ* haplotypes. Edges are colored based on their loop structure (CspZ_B379_, CspZ_B408_, and CspZ_B31_ in green, orange, and purple, respectively).

**Figure S10.**
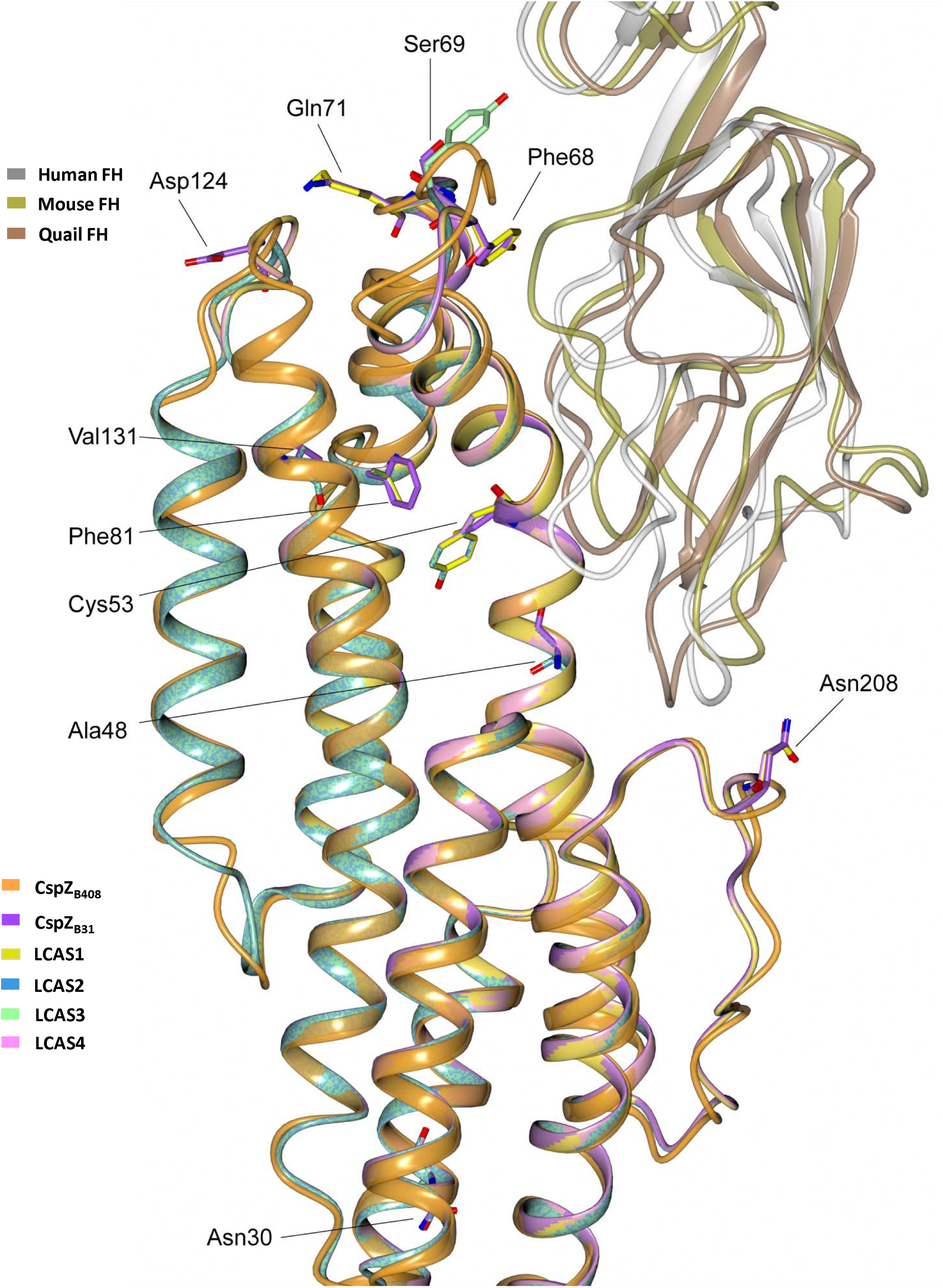
The CspZ last common ancestor states are predicted to versatiley bind FH. The crystal structure of CspZ_B408_-SCR6-7 where human SCR6-7 (grey) is superimposed with mouse FH SCR6-7 (gold, PDB: 2YBY) and the predicted structure of quail SCR6-7 (brown). CspZ_B408_ (orange) is superimposed with CspZ_B31_ (purple) and the last common ancestor states: LCAS1 (yellow), LCAS2 (blue), LCAS3 (green), and LCAS4 (pink). Residues that differ between CspZ_B31_ and the LCAS variants are labelled.

**Figure S11.**
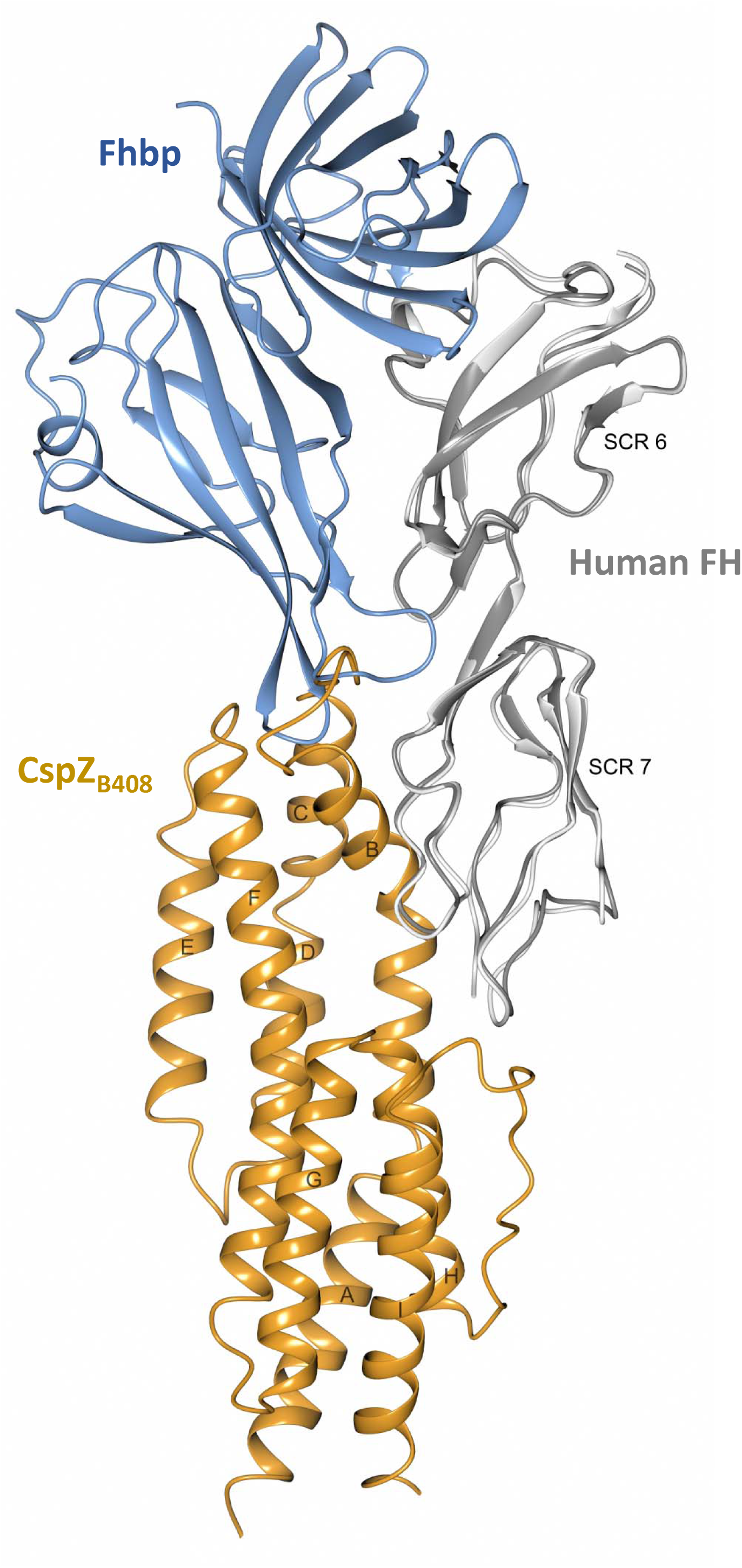
Structural comparison between CspZ_B408_-human SCR6-7 and the *N. meningitidis* Fhbp-human SCR6-7 complexes. Human FH SCR6-7 (light grey) from the complex structure with CspZ_B408_ (orange) was superimposed with human FH SCR6-7 (dark grey) from the complex structure with Fhbp (dark blue, PDB: 2W81), the FH-binding protein from *N. meningitidis.* α- helices in CspZ_B408_ are labelled from A to I starting from the N-terminus.

**Figure S12.**
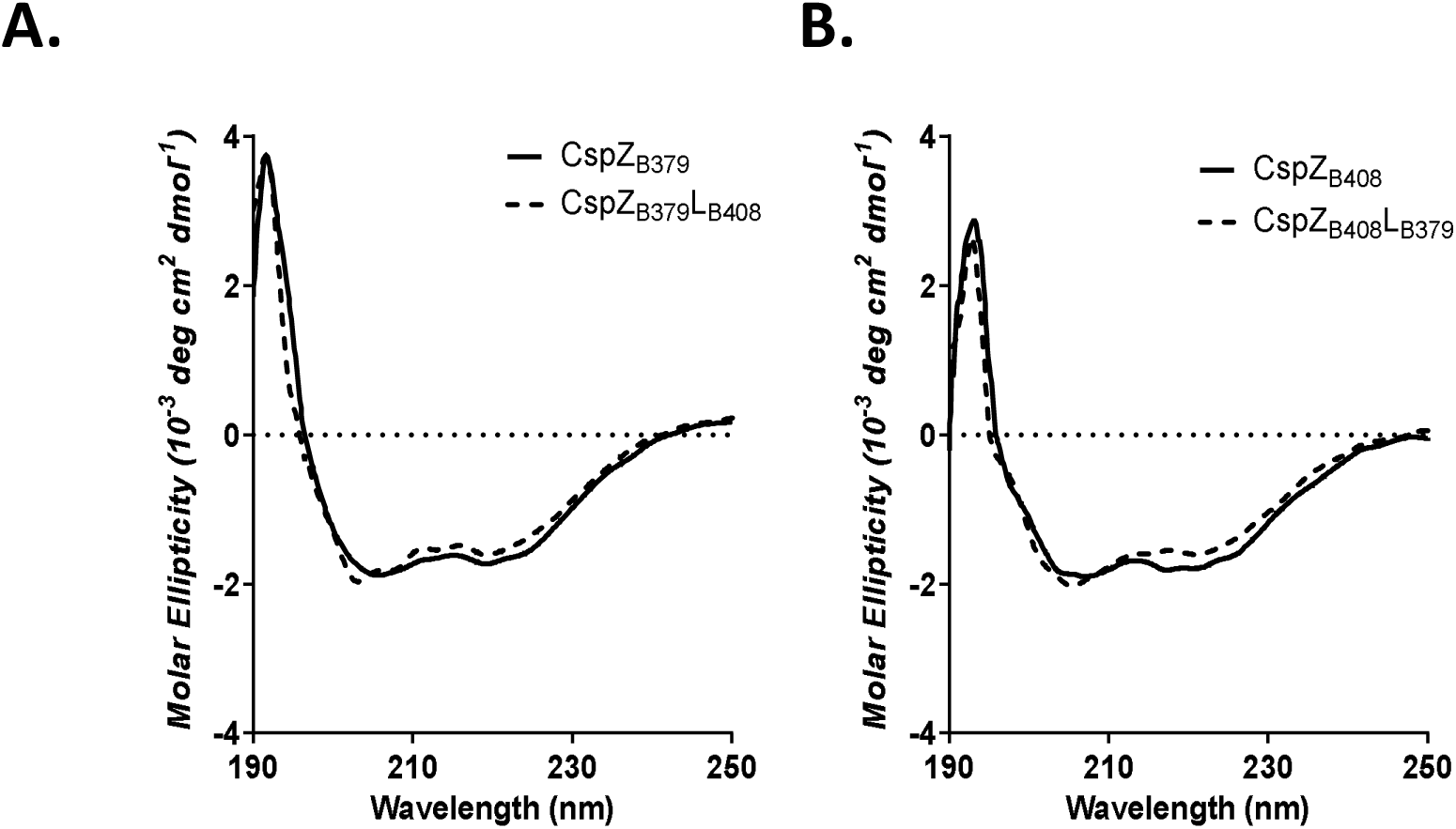
CD spectra demonstrate no impacts of secondary structures by swapping the loops. Far-UV CD analysis of CspZ_B379_, CspZ_B408_, CspZ_B379_L_B408_, and CspZ_B408_L_B379_. The molar ellipticity, Φ, was measured from 190 to 250nm for 10μM of each protein in PBS.

**Figure S13.**
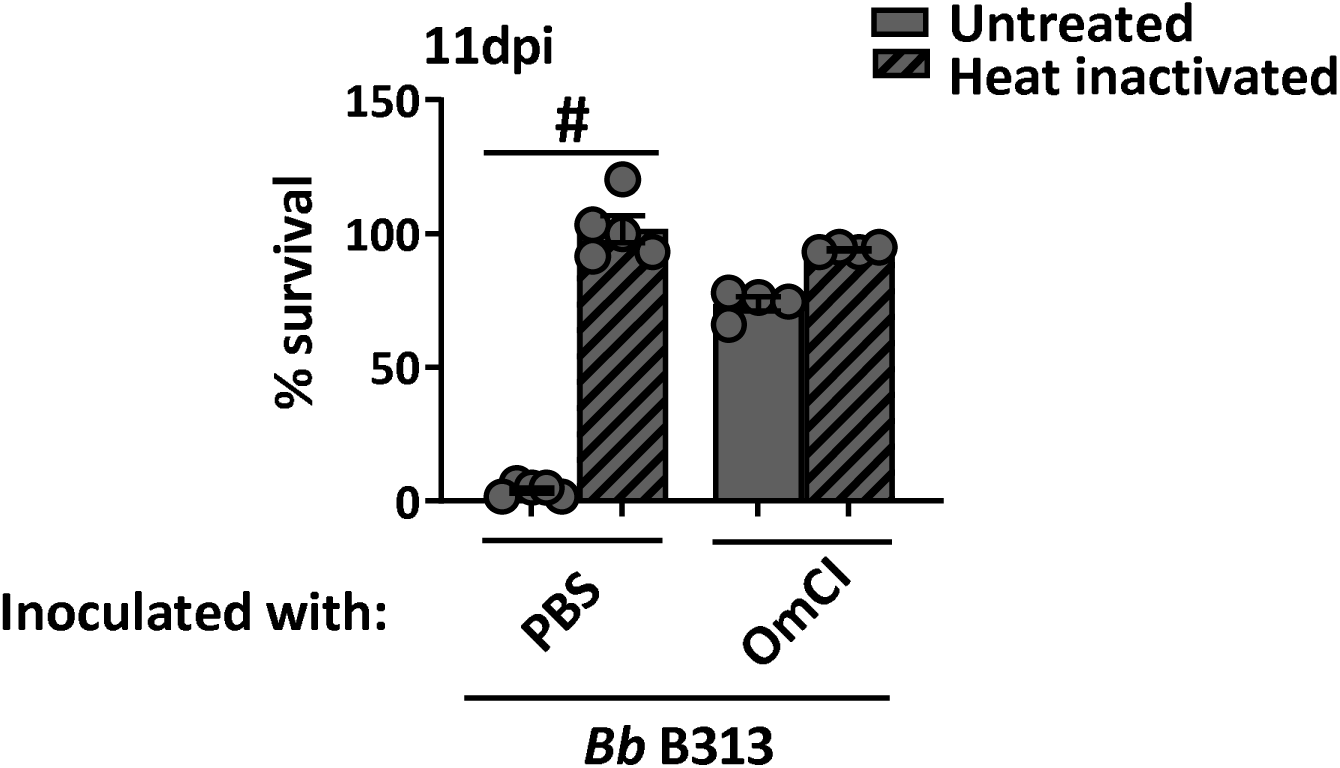
OmCI prevents quail serum-mediated killing of a complement-sensitive spirochete strain at 11 days post injection. *Coturnix* quail were subcutaneously inoculated with OmCI (1mg/kg of quail) or PBS buffer. Untreated (filled bars) or heat-treated (hatched bars) sera collected from these quail at 11 days post inoculation (dpi) were incubated with a serum-sensitive, highly passaged *B. burgdorferi* strain B313. The number of motile spirochetes were quantified microscopically and the survival percentage of the spirochetes was calculated using the number of mobile spirochetes at 4 h post incubation normalized to that at 0 h. Each bar represents the mean ± SEM of three independent experiments from sera from four quail per group. Significant differences (p < 0.05, Mann-Whitney test) in the percentage survival of spirochetes are indicated (“#”).

## SUPPLEMENTAL TABLES

**Table S1:** The list of CspZ_B408_ residues that binds to human FH based on CspZB408-human SCR6-7 complex structure and their equivalent residues of CspZ_B31_ and CspZ_B379_.

| **CspZ<sub>B408</sub>** | **CspZ<sub>B31</sub>** | **CspZ<sub>B379</sub>** |
| --- | --- | --- |
| Asp47 | Asp47 | Asp47 |
| Tyr50 | Tyr50 | Tyr50 |
| Asn51 | Asn51 | Ser51 |
| Thr54 | Thr54 | Thr54 |
| Asn58 | Asn58 | Asn58 |
| Thr62 | Thr62 | Thr62 |
| Asp71 | - | - |
| Asp73<sup>a</sup> | Asp70<sup>a</sup> | Asp74<sup>a</sup> |
| Ser75<sup>a</sup> | Ser72<sup>a</sup> | Ser76<sup>a</sup> |
| Arg142 | Arg139 | Arg143 |
| Asn183 | Asn180 | Asn184 |
| Tyr214 | Tyr211 | Tyr215 |

<sup>a</sup>Interacts with SCR6.

**Table S2.** CspZ variants differ in binding to Factor H from different animals.

| CspZ variant | Factor H source | ELISA<sup>a</sup> | ----- Surface Plasmon Resonance<sup>b</sup> ----- |  |  |
| --- | --- | --- | --- | --- | --- |
|  |  | K<sub>D</sub> (μM) | K<sub>D</sub> (μM) | k<sub>on</sub> (10<sup>3</sup>s<sup>-1</sup>M<sup>-1</sup>) | k<sub>off</sub> (s<sup>-1</sup>) |
| CspZ<sub>B31</sub> | Mouse | 0.43±0.74<sup>c</sup> | 0.20±0.02 | 45.78±15.36 | 0.0084±0.0022 |
|  | Quail | 0.91±0.08<sup>d</sup> | 0.81±0.01 | 117.43±3.25 | 0.095±0.0032 |
| CspZ<sub>B379</sub> | Mouse | n.b.<sup>d</sup> | n.b. | n.b. | n.b. |
|  | Quail | 0.59±0.037 | 0.75±0.15 | 119.83±7.08 | 0.088±0.012 |
| CspZ<sub>B379</sub>L<sub>B408</sub> | Mouse | 1.39±0.12 | 0.90±0.12 | 21.16±10.58 | 0.018±0.0094 |
|  | Quail | n.b. | n.b. | n.b. | n.b. |
| CspZ<sub>B408</sub> | Mouse | 0.68±0.05 | 0.20±0.02 | 22.13±1.31 | 0.008±0.002 |
|  | Quail | n.b. | n.b. | n.b. | n.b. |
| CspZ<sub>B408</sub>L<sub>B379</sub> | Mouse | n.b. | n.b. | n.b. | n.b. |
|  | Quail | 1.38±0.47 | 0.99±0.06 | 72.43±2.14 | 0.072±0.006 |
| GST<sup>e</sup> | Mouse | n.b. | n.d. | n.d. | n.d. |
|  | Quail | n.b. | n.d. | n.d. | n.d. |

All values represent the mean ± SEM of three experiments.
<sup>a</sup>Determined using GST tagged CspZ variants or mutant proteins.
<sup>b</sup>Determined using untagged CspZ variants or mutant proteins.
<sup>c</sup>Reported previously in (41).
<sup>d</sup>No binding activity was detected.
<sup>e</sup>GST was included as a negative control.

**Table S3.**
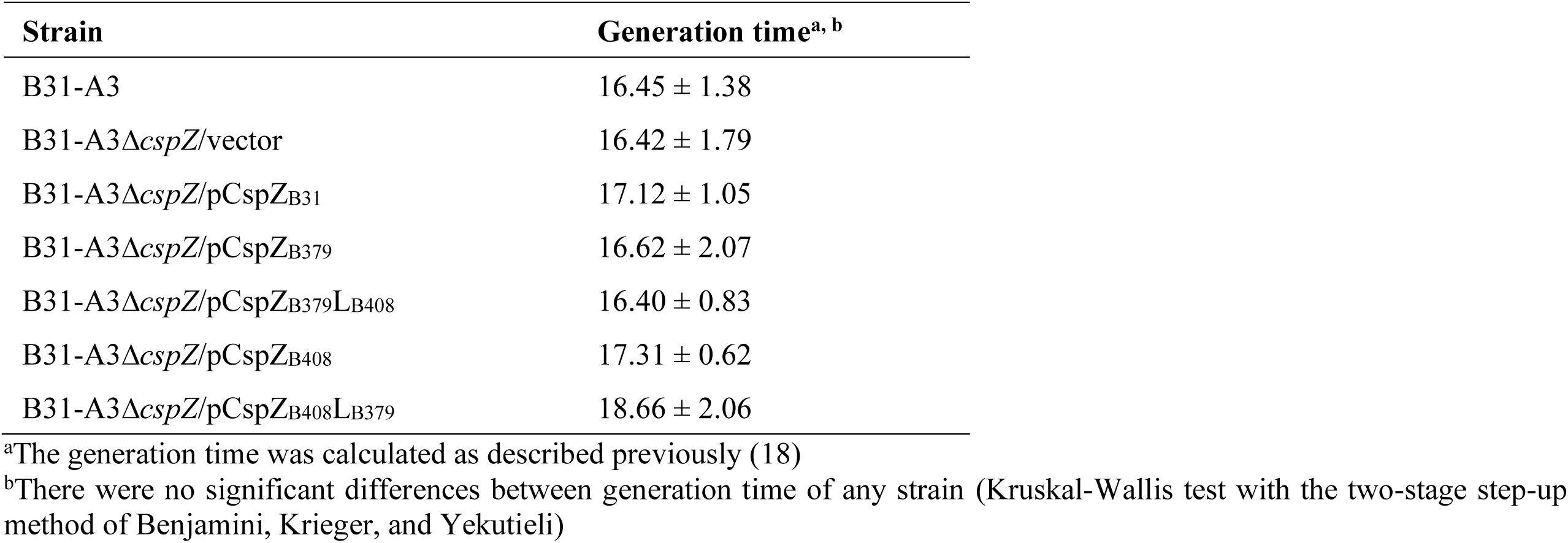
The generation time for *B. burgdorferi* strains used in this study.

**Table S4:**
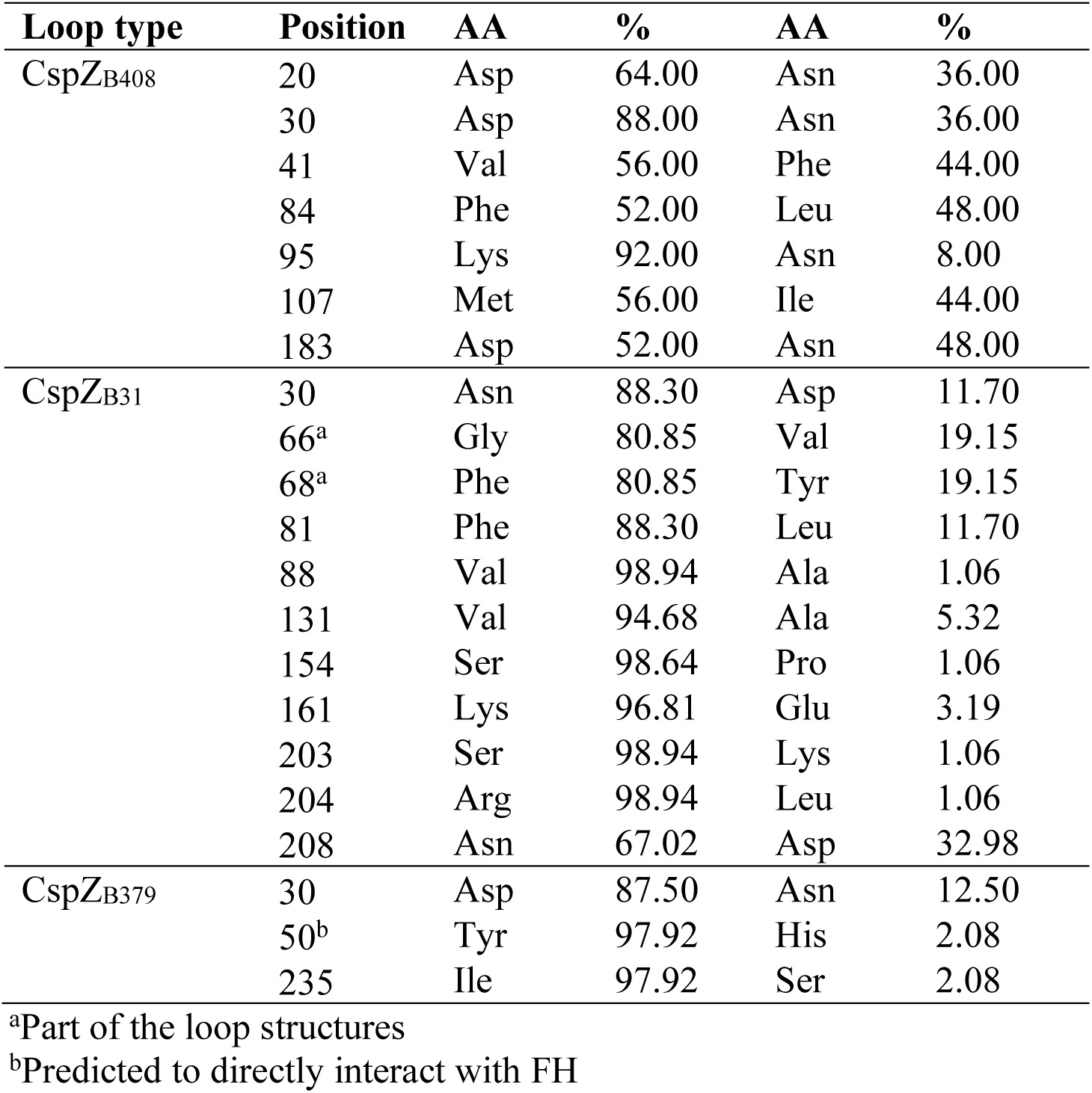
The percentage of SAPs in CspZ variants.

| Loop type | Position | AA | % | AA | % |
| --- | --- | --- | --- | --- | --- |
| CspZ<sub>B408</sub> | 20 | Asp | 64.00 | Asn | 36.00 |
|  | 30 | Asp | 88.00 | Asn | 36.00 |
|  | 41 | Val | 56.00 | Phe | 44.00 |
|  | 84 | Phe | 52.00 | Leu | 48.00 |
|  | 95 | Lys | 92.00 | Asn | 8.00 |
|  | 107 | Met | 56.00 | Ile | 44.00 |
|  | 183 | Asp | 52.00 | Asn | 48.00 |
| CspZ<sub>B31</sub> | 30 | Asn | 88.30 | Asp | 11.70 |
|  | 66<sup>a</sup> | Gly | 80.85 | Val | 19.15 |
|  | 68<sup>a</sup> | Phe | 80.85 | Tyr | 19.15 |
|  | 81 | Phe | 88.30 | Leu | 11.70 |
|  | 88 | Val | 98.94 | Ala | 1.06 |
|  | 131 | Val | 94.68 | Ala | 5.32 |
|  | 154 | Ser | 98.64 | Pro | 1.06 |
|  | 161 | Lys | 96.81 | Glu | 3.19 |
|  | 203 | Ser | 98.94 | Lys | 1.06 |
|  | 204 | Arg | 98.94 | Leu | 1.06 |
|  | 208 | Asn | 67.02 | Asp | 32.98 |
| CspZ<sub>B379</sub> | 30 | Asp | 87.50 | Asn | 12.50 |
|  | 50<sup>b</sup> | Tyr | 97.92 | His | 2.08 |
|  | 235 | Ile | 97.92 | Ser | 2.08 |

<sup>a</sup>Part of the loop structures
<sup>b</sup>Predicted to directly interact with FH

**Table S5.** Estimated diversification times for each lineage.

| **Lineage** | **Median<sup>a</sup>** | **HPD<sub>5%</sub><sup>a</sup>** | **HPD<sub>95%</sub><sup>a</sup>** |
| --- | --- | --- | --- |
| CspZ-B31 | 784 | 263 | 1860 |
| CspZ-B379 | 261 | 59 | 741 |
| CspZ-B408 | 671 | 179 | 1679 |

<sup>a</sup>Years before present

**Table S6.**
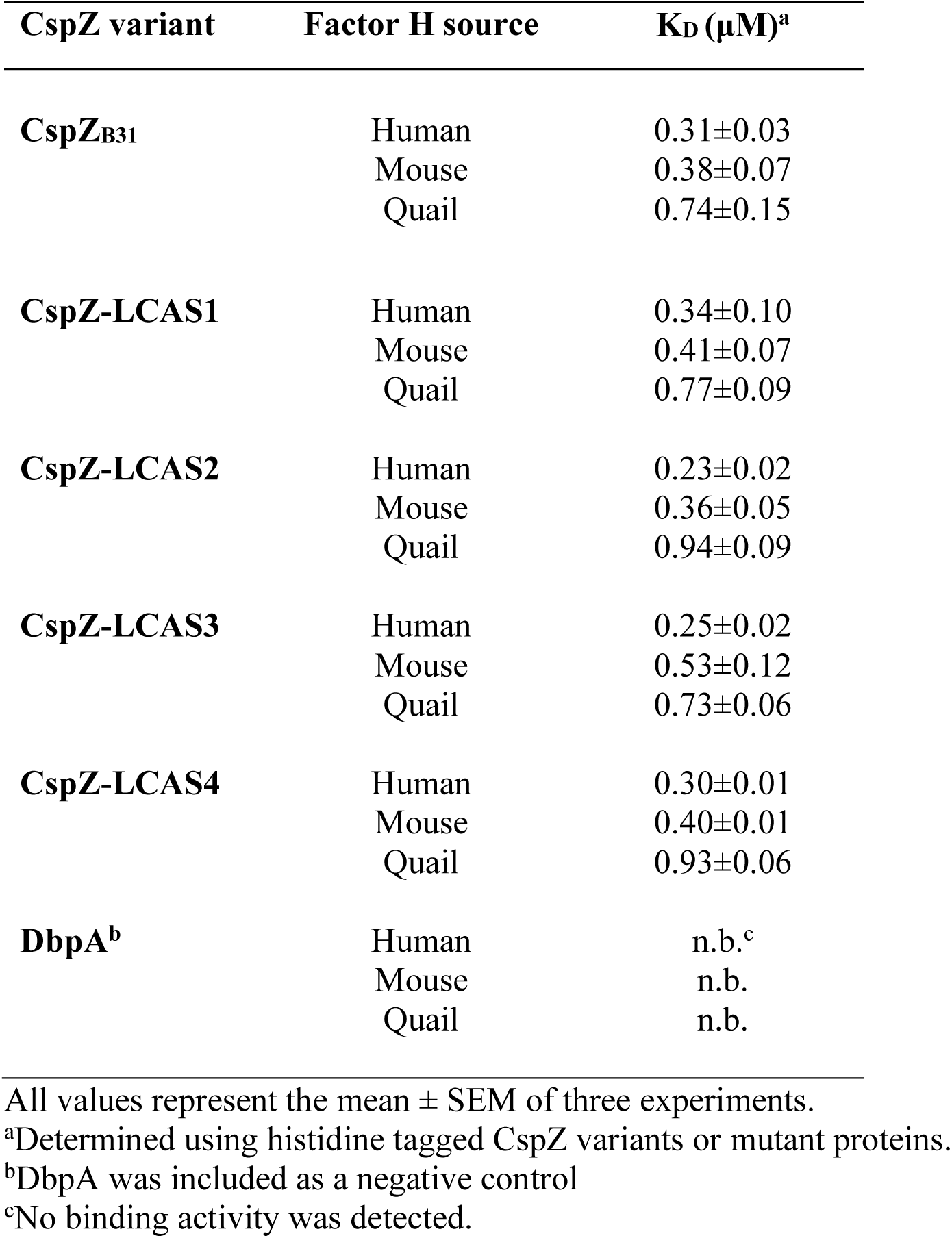
CspZ variants of last common ancestral states display versatile binding ability to human, mouse, and quail FH.

**Table S7.**
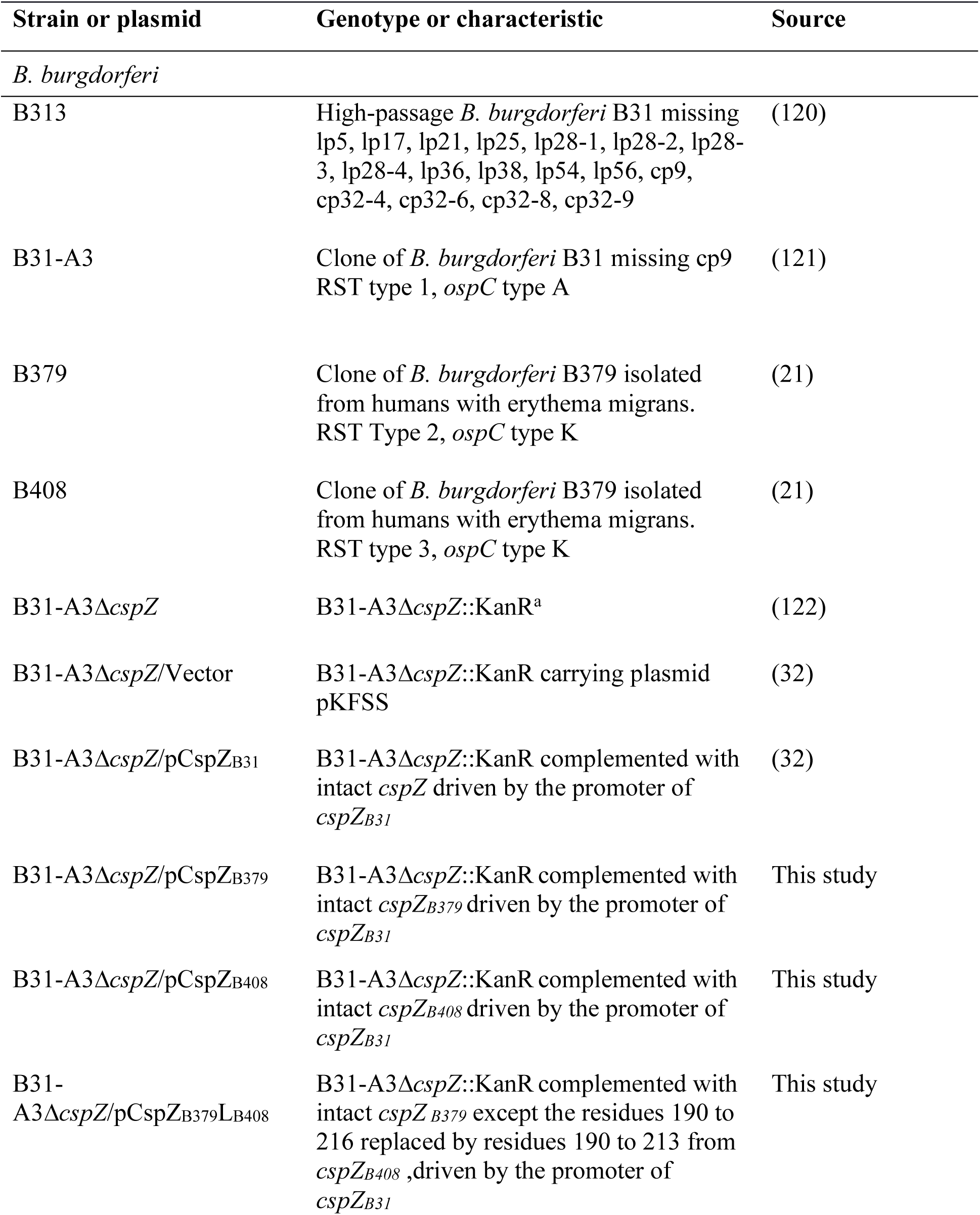
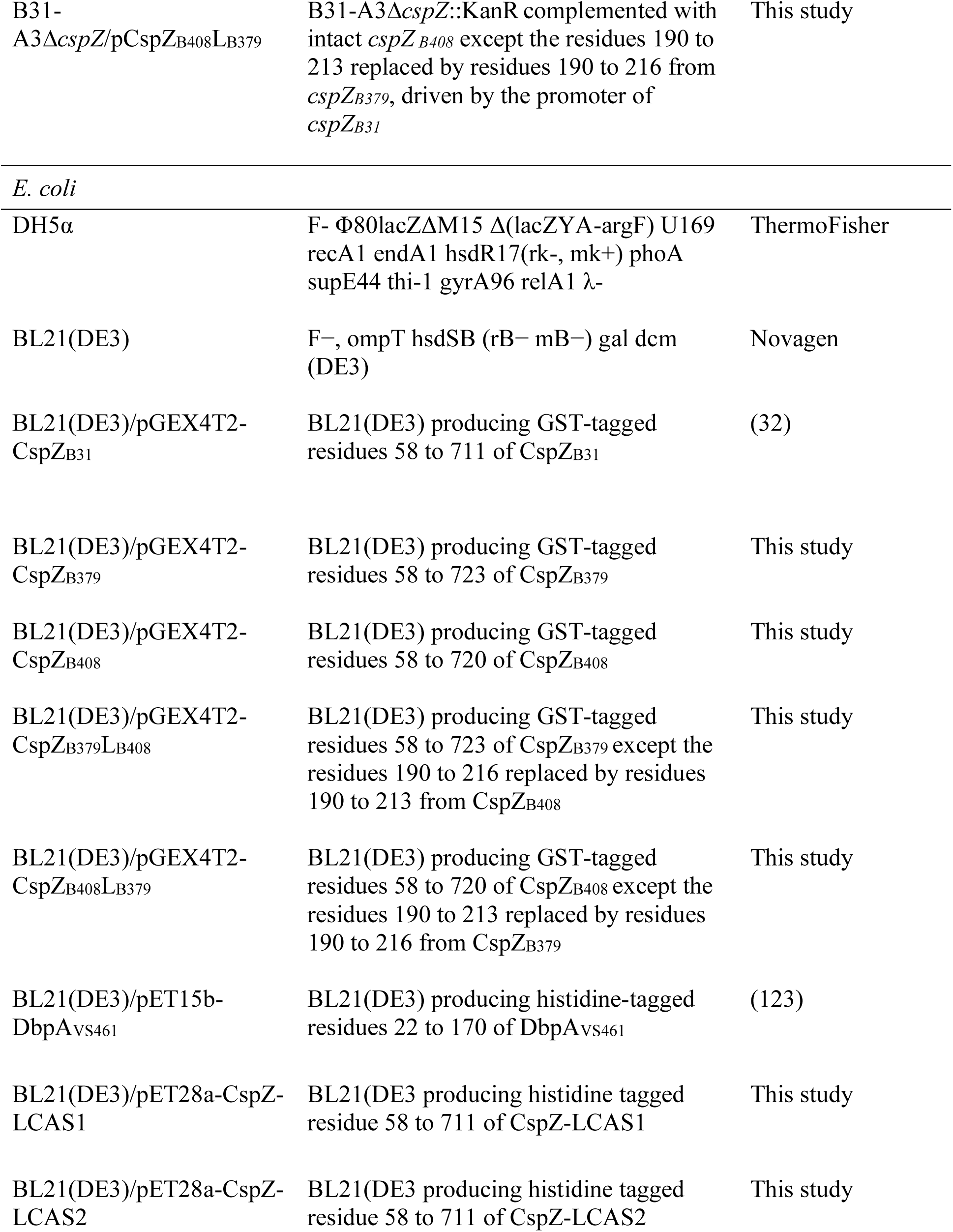
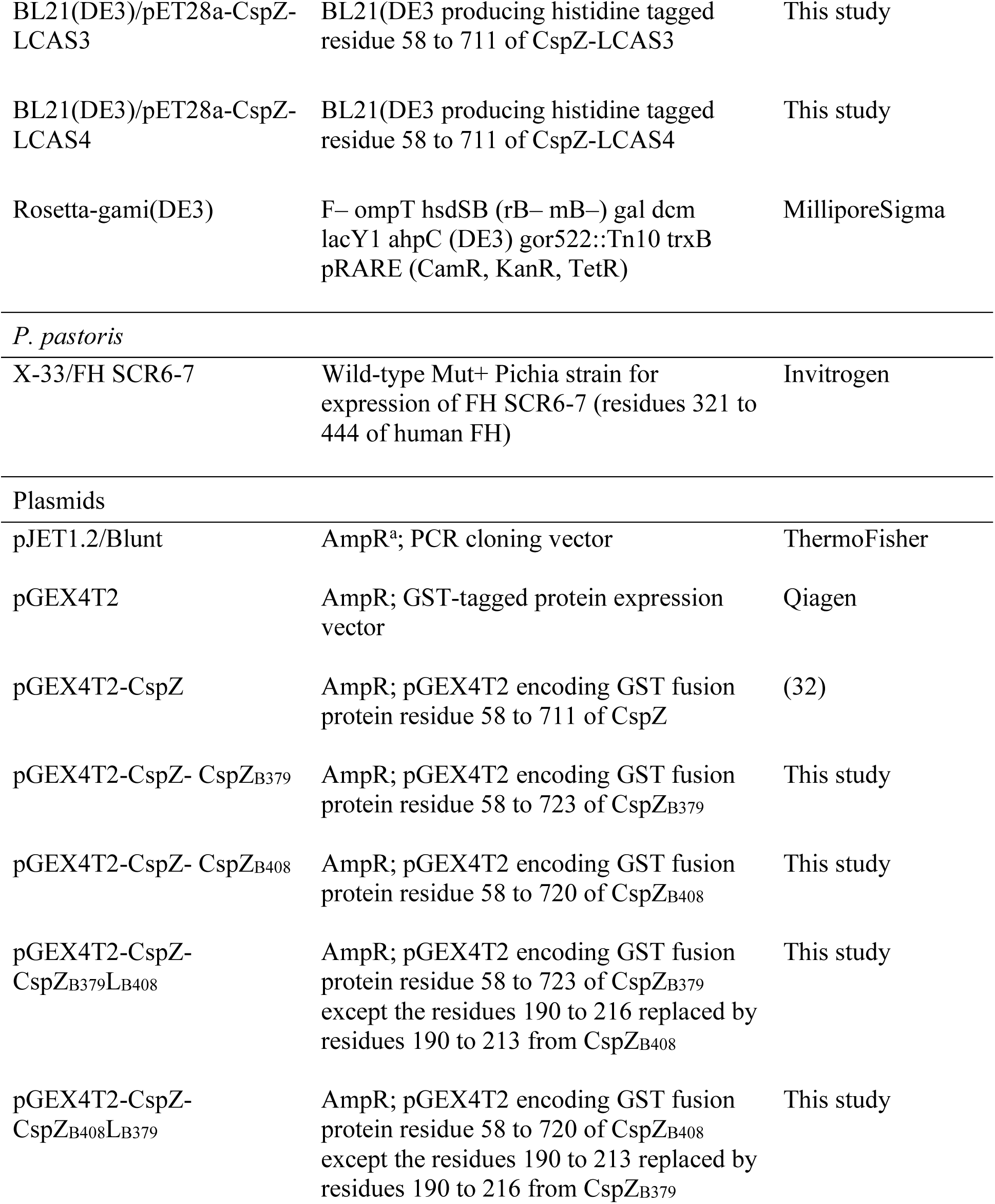
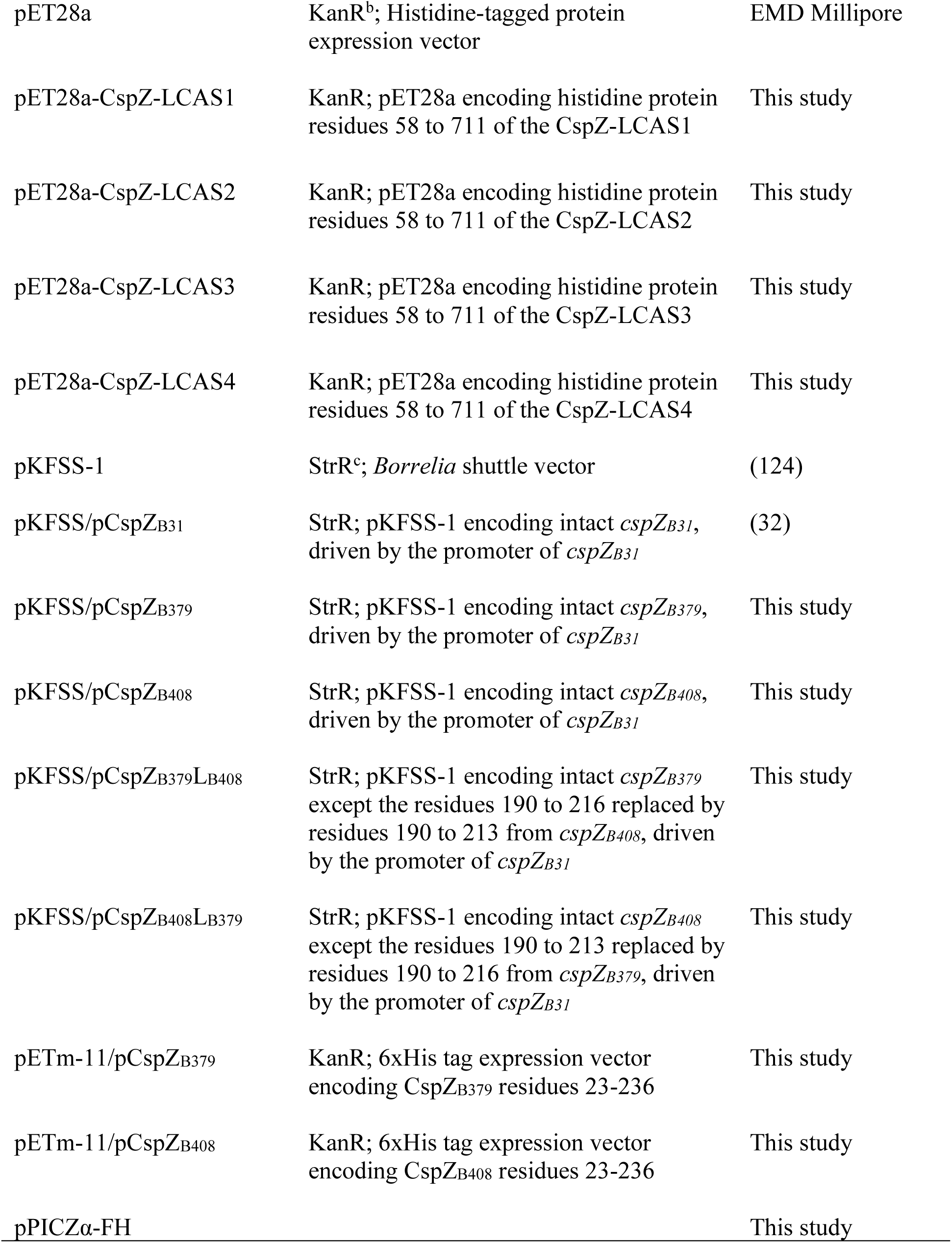
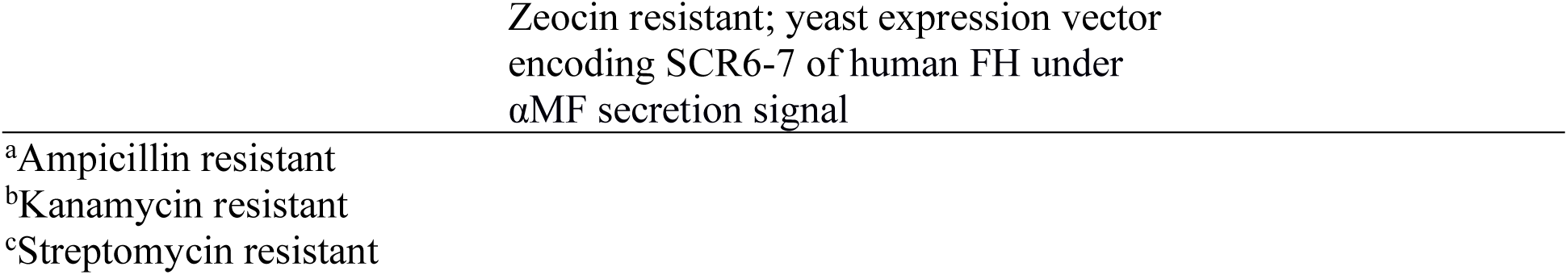
The strains and plasmids used in this study.

**Table S8.**
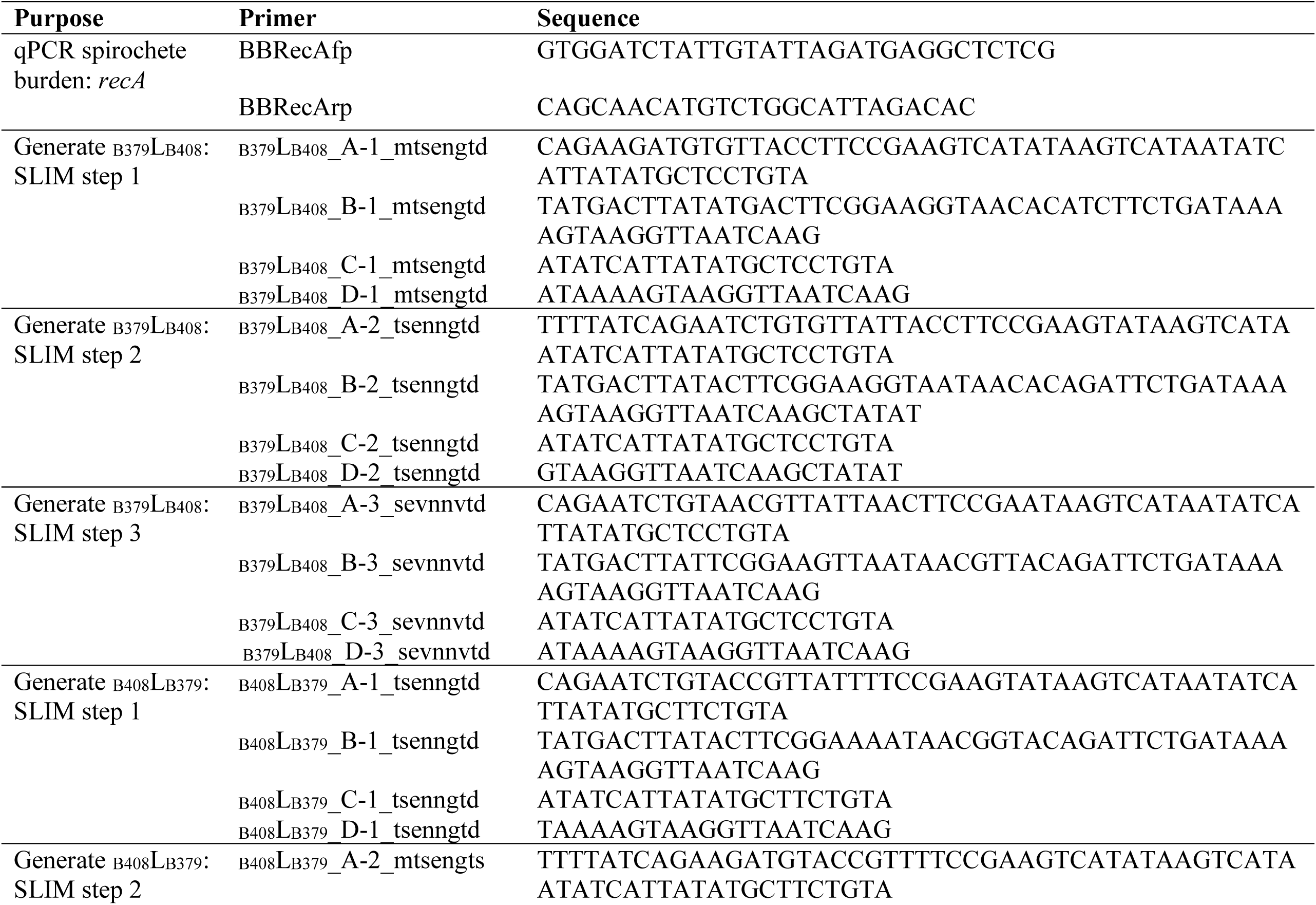
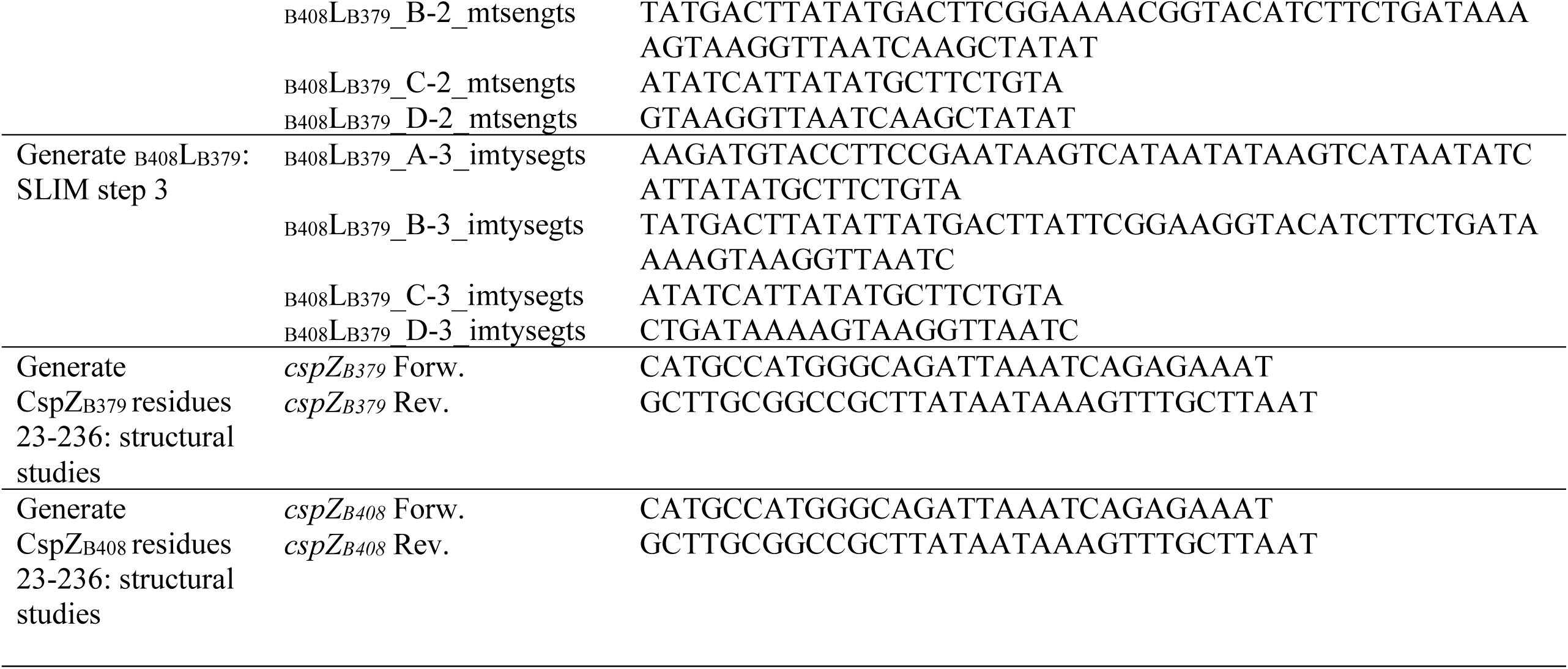
Primers used in this study.

**Table S9.**
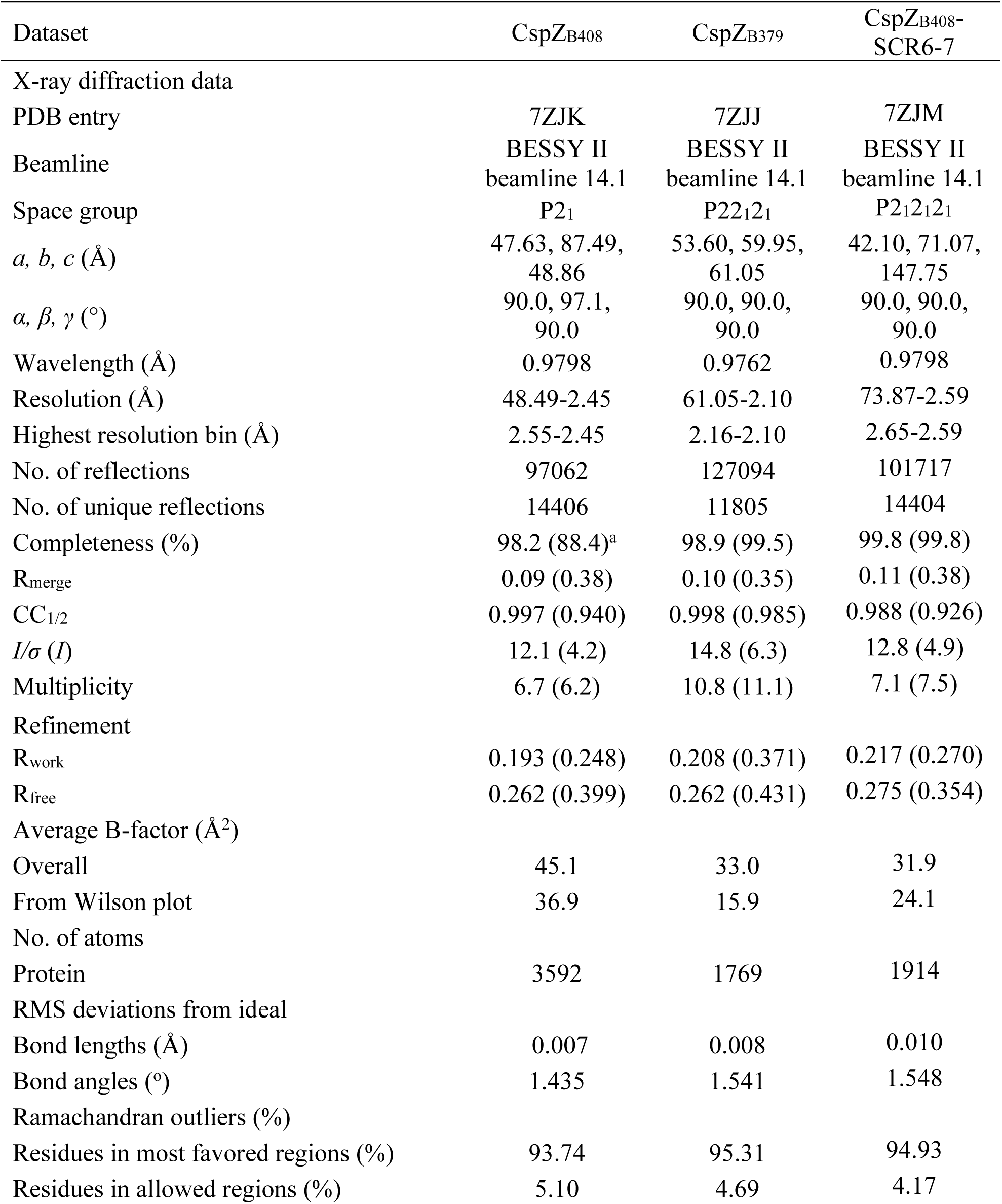
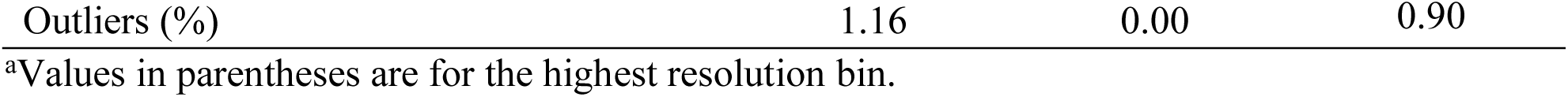
Data processing, refinement, and validation statistics of crystal structures.

## Notes

### Competing Interest Statement

The authors have declared no competing interest.

### Summary of Updates

Acknowledgements

